# The stereochemical mechanism of the B_12_-dependent radical SAM glutamine methyltransferase (QCMT): Novel insights and unprecedented post-translational modifications

**DOI:** 10.64898/2026.03.17.712072

**Authors:** Thibaut Bourdin, Alain Guillot, Mickaël Mauger, Benjamin Lefranc, Sylvain Gervason, Matthieu Glousieau, Stéphane Grimaldi, Jérôme Leprince, Aurélien Thureau, Alhosna Benjdia, Olivier Berteau

## Abstract

Methyl-coenzyme M reductase (MCR) is a crucial enzyme for methanogenesis and harbors several unusual post-translational modifications. Recent studies have identified glutamine C-methyltransferase (QCMT), as a B_12_-dependent radical SAM enzyme responsible for methylating a glutamine residue within the MCR active site. B_12_-dependent radical SAM enzymes have the remarkable ability to alkylate unactivated C*sp^2^*- and *Csp^3^*-atoms in a stereoselective manner. However, the factors influencing the stereo-selectivity and catalytic properties of this emerging superfamily of enzymes remain poorly understood. In this study, we report the mechanistic, structural, and biochemical investigation of several QCMTs. Our findings reveal significant differences among them, notably in their ability to bind cobalamin. In addition, our data support that Cα H-atom abstraction and methyl transfer are not concerted but rather independent processes that require motion within the enzyme’s active site. We also demonstrate that QCMT can catalyze novel reactions, including the formation of unnatural *C*-methylated residues, peptide epimerization, reversible H-atom abstraction, and the direct conversion of glycine into d-alanine. Overall, our data are consistent with QCMT being a unique and versatile biocatalyst allowing for the installation of unnatural post-translational modifications and provide a structural and biochemical rationale for the control of the stereochemistry by B_12_-dependent radical SAM enzymes.

## Introduction

Radical SAM (*S*-adenosylmethionine) enzymes, with over 800,000 distinct proteins identified, are considered the most versatile and widespread biocatalysts^1–6^. These enzymes, among other transformations^3,7–9^, catalyze a wide range of post-translational modifications, from thioether bond formation^10,11^ and radical activation^9,12–14^ to anaerobic oxidation^15,16^ and methylthiolation^17–19^. Radical SAM enzymes have evolved to incorporate additional domains, expanding their catalytic repertoire. Among them, with more than 220,000 members, the vitamin B_12_-dependent radical SAM enzymes^1^ built on a radical SAM and a vitamin B_12_ (cobalamin) binding domain, form the largest class^1,5,20^. These biocatalysts possess the remarkable ability to alkylate various unreactive C*sp^2^*- (*e.g.,* TsrM^21,22^) and C*sp^3^*-hybridized atoms (*e.g.,* TokK^23^, PoyC^24^, CysS^25^, Mmp10^5,21,26–28^), in addition to performing several unrelated reactions^1^. Initially, it was proposed that these enzymes contained an *N*-terminal B_12_-binding domain and a *C*-terminal radical SAM domain^5,26,29^. However, recent studies revealed a more complex picture^1,5,26,30^.

While B_12_-dependent radical SAM enzymes can modify a broad range of substrates, from secondary metabolites to proteins, they are believed to utilize a conserved radical mechanism. Many of these enzymes invert the stereochemistry of the methylated carbon atom, in agreement with the unique architecture of their active site, which features both an activation center and an alkylation center surrounding the substrate. Conversely, some enzymes retain the stereochemistry of the substrate. For example, based on the structure of its product, (*S*)-2-methylglutamine (Me-Gln), QCMT which methylates a glutamine residue within the active site of the alpha-subunit of a subset of methyl-coenzyme M reductases (MCR)^26,31–33^, the central enzyme in methane production in methanogenic archaea^34–36^, is proposed to retain the stereochemistry of the methylated carbon atom^37–39^. Interestingly, Me-Gln is not strictly conserved in MCR, whereas other post-translational modifications, such as Me-Arg (installed by the methanogenesis marker protein 10, Mmp10^26,31^), are universal^26,31,40–43^. The exact role of these post-translational modifications remains unclear, but they are believed to be important for optimal *in vivo* activity of MCR and thus for methane metabolism^34^.

Currently, approximately half of the experimentally investigated B_12_-dependent radical SAM enzymes such as MoeK^44^, GenD1^45^, Fom3^46^ and TokK^47^, have been shown to invert the stereochemistry of their substrates, while others like GenK^48^, Orf29^49^ and QCMT, retain it.

In the last couple of years, several research groups have focused on QCMT, demonstrating its biological function and elucidating its crystallographic structure^32,50^. However, no rationale has been provided for how this enzyme controls the stereochemistry of its reaction. To gain further insights into this central question and into B_12_-dependent radical SAM enzymes in general, we undertook the biochemical and structural characterizations of several QCMT enzymes. Unexpectedly, we discovered that QCMT, owing to its unique mechanism, can catalyze novel reactions, including reversible hydrogen abstraction, peptide epimerization, and unnatural alkylation reactions, including the direct conversion of glycine into d-alanine. Our study demonstrates the unique potential of this enzyme for applications in synthetic biology, late-stage labeling, and peptide modification. In addition, structural investigation supports the notion that cobalamin binding induces a conformational change and that, following substrate activation, the transfer of the methyl group is likely an independent process, with the Cα hydrogen abstraction being a partially rate-determining step. Finally, we also illustrate the biocatalytic potential of QCMT by showcasing its ability to catalyze unnatural post-translational modifications in peptides, thereby expanding the chemical repertoire of radical SAM enzymes.

### QCMT Efficiently Methylates Short MCR Mimics

To investigate the mechanism and substrate promiscuity of QCMT, we cloned several homologs from *Methanoculleus thermophilus*, *Methanothermococcus thermolithotrophicus*, *Methanoculleus marisnigri*, *Methanothermococcus okinawensis* and *Methanoculleus bourgensis* as strep-TAG fusion proteins and expressed them in *E. coli*. The most active and soluble ones, hereafter referred as **QCMT_1_** and **QCMT_2_**, were the ones from *M. thermophilus* and *M. thermolithotrophicus* recently identified by the Layer Lab^31,32^ (see **Supplementary Fig. S1**). They were thus selected for further biochemical and structural characterization. Analysis by UV-visible spectroscopy and LC-MS revealed that while **QCMT_1_**binds cobalamin as previously reported^32^, **QCMT_2_** only marginally binds cobalamin (**Supplementary Fig. S1**). Further characterization of **QCMT_1_**, which can be isolated in both B_12_-bound and free forms, was performed by anaerobic size-exclusion chromatography coupled with small-angle X-ray Scattering (SEC-SAXS). As shown, the SEC-SAXS profile of **QCMT_1_** indicated that it is a monomeric, homogeneous, globular protein, with a radius of gyration (Rg) of 24.35 ±0.04 Å and an estimated molecular weight of 41 kDa (**Fig. 1**). Incubation with cobalamin resulted in a significant and reproducible decrease in the Rg (Rg = 24.03 ±0.04 Å), indicating a conformational change in the protein. Conversely, the addition of SAM alone, or in the presence of the peptide substrate, did not cause any measurable structural changes. Overall, the SEC-SAXS analysis suggests that cobalamin induces small but significant conformational changes upon binding, leading to protein compaction, a change that was not observed in OxsB, the only B12-dependent radical SAM enzyme (albeit not a methyltransferase) structurally characterized in the absence of cobalamin^51^.

**Figure 1.**
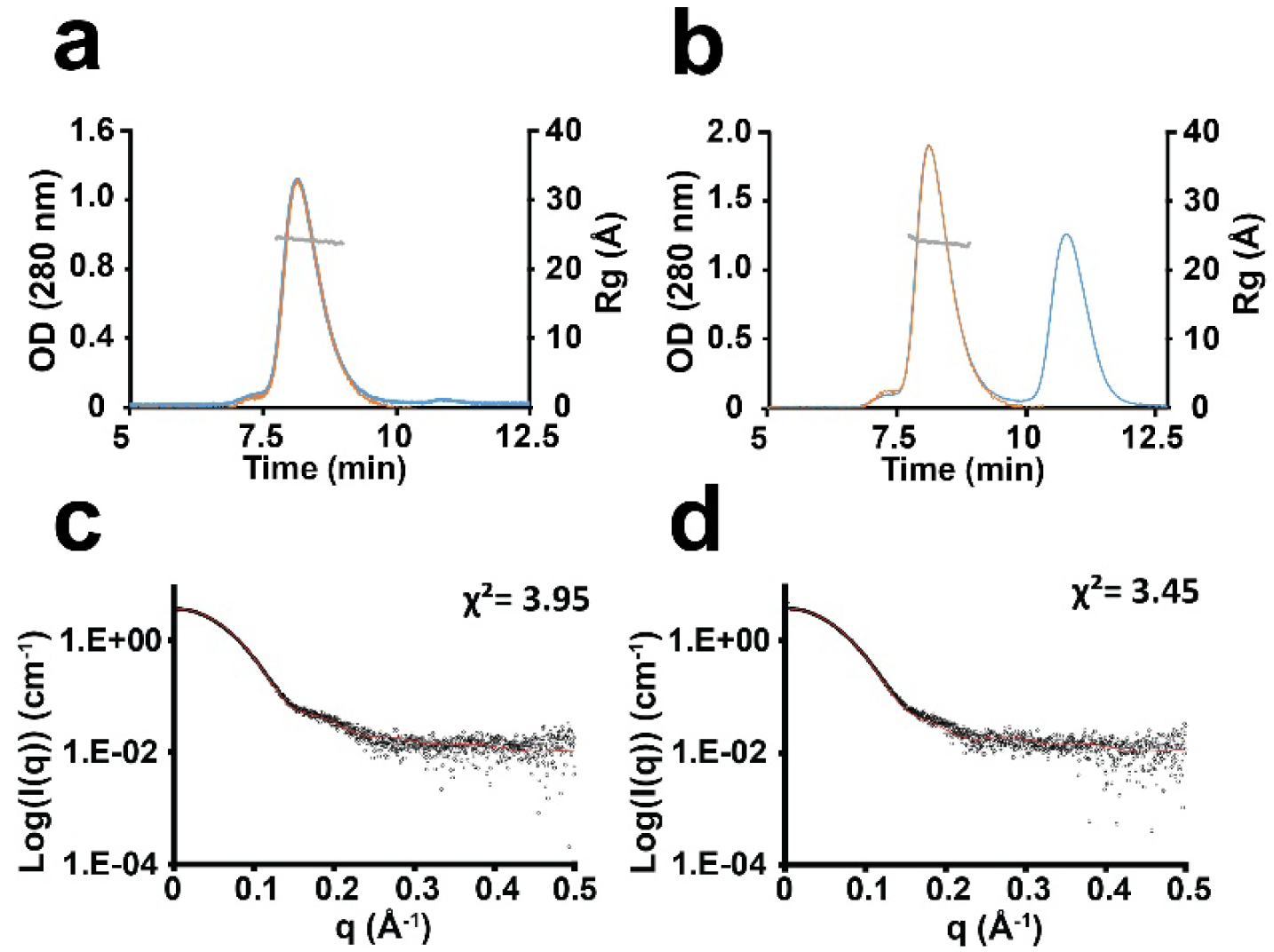
SEC-SAXS analysis of QCMT_1_ in the absence (a) or the presence (b) of OHCbl. The UV–visible signal at λ = 280 nm (blue) and the SAXS intensity at zero angle (orange) are superimposed with the computed Rg from SAXS data (gray circles). The peak eluting at ∼11 min corresponds to free OHCbl. SAXS curves of **QCMT_1_** in the absence (**c**) or the presence (**d**) of OHCbl. Fit curves using PepsiSAXS^52^ are indicated by a red trace and χ^2^ value determined between the experimental data and the simulated model (**Supplementary Table S1**).

In previous studies^32,50^, **QCMT_1_**was tested with a 24-mer peptide; however, the size and hydrophobicity of this substrate likely hindered the quantification of all the reaction products formed. Indeed, according to its proposed mechanism (**Fig. 2a**), QCMT produces both 5’-dA and SAH (resulting from radical cleavage and alkylation mediated by the SAM cofactor, respectively), as well as the methylated peptide/protein. To address this issue, we hypothesized that a shorter synthetic peptide (13-mer) framing the targeted glutamine residue would serve as a better substrate. Indeed, a 13-mer peptide has been shown to be sufficient to interact with Mmp10, the only B_12_-dependent radical SAM enzyme structurally characterized in complex with its peptide substrate^26^. Both **QCMT_1_** and **QCMT_2_**were reconstituted using standard protocols in order to obtain holoproteins with their intact [4Fe-4S] clusters. They were assayed under anaerobic conditions, in the presence of hydroxycobalamin (OHCbl), SAM, peptide **1Q** (13-mer, [M+3H]³⁺: 438.89), and titanium (III) citrate (Ti(III)), as a one-electron donor. As shown, we obtained efficient conversion of **1Q** into a methylated product, **CH_3_-1Q** ([M+3H]³⁺: 443.56). The location of the methyl group on Gln-7 (corresponding to Gln403 in the α-subunit of MCR from *M. thermolithotrophicus*) was unequivocally confirmed by LC-MS/MS (**Fig. 2b**, **c**, and **Supplementary Fig. S2**).

**Figure 2.**
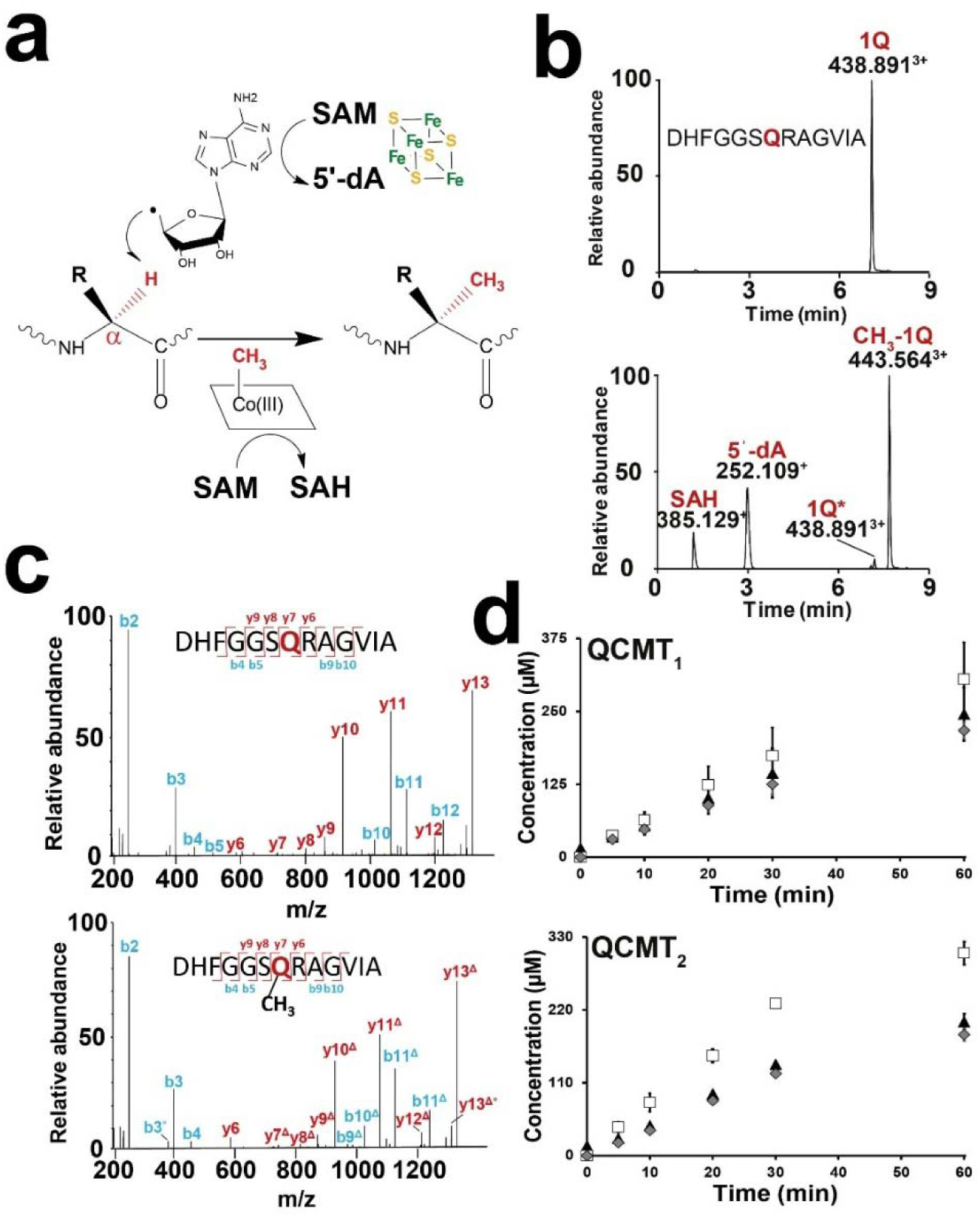
QCMT biochemical assay on a short synthetic peptide. (**a**) Mechanistic hypothesis for QCMT alkylation reaction. One molecule of SAM is reductively cleaved for H-atom abstraction leading to the generation of 5′-dA, while a second molecule of SAM is used for MeCbl regeneration. R: Amino acid side chain (**b**) LC-MS analysis before (upper trace) and after (lower trace) 30 min incubation of peptide **1Q** with **QCMT_1_**. (**c**) MS fragmentation of peptide **1Q** (upper trace) and **CH_3_-1Q** (lower trace). Ions with a +14.015 Da addition mass corresponding to an addition of a CH_2_ are noted by Δ. See **Supplementary Fig. S2** and **Tables S2 & S4** for full assignment. (**d**) Time course analysis of the reaction catalyzed by **QCMT_1&2_**: SAH (triangle symbols), 5’-dA (diamond symbols) and **CH_3_-1Q** (square symbols). Experiments were performed in triplicate. **QCMT** (7.5 µM) was incubated after iron-sulfur reconstitution with OHCbl (0.1 mM), SAM (0.5 mM) and **1Q** (150 µM) (panels **b** and **c**) or with MeCbl (0.1 mM), SAM (0.8 mM) and **1Q** (300 µM) (panel **d**), under anaerobic and reducing conditions. Reactions were performed at 50°C and initiated by adding titanium (III) citrate (2 mM).

To quantify all the products formed, we developed an LC-MS method and optimized the reaction conditions. Notably, we monitored the effects of reactants’ concentration and incubation temperature. Both enzymes showed enhanced activity at elevated temperatures with maximum activity at 70°C and 50°C for **QCMT_1_** and **QCMT_2_**, respectively (**Supplementary Fig. S3**). However, at temperatures >50°C, significant 5′-dA and SAH degradation was evidenced, and the activity of **QCMT_2_** was impaired. The temperature of 50°C was thus chosen for the quantification assays.

Under these assay conditions, we determined specific activities of 167.2 ±42.6 µmol·min⁻¹·mg⁻¹ for **QCMT_1_** and 193.8 ±2.7 µmol·min⁻¹·mg⁻¹ for **QCMT_2_**, for the formation of the methylated product (**CH_3_-1Q**) (**Fig. 2d**). With a *kcat* value of approximately ∼0.7 min⁻¹ for methylated peptide, both enzymes exhibited an activity that was 10 to 20 times faster than the one previously reported^32^ and an apparent turnover of ∼40 in 1 hour (**Fig. 2d**). It should be noted that this turnover is likely underestimated, since a large portion of the substrate was converted by the end of the reaction.

In contrast to recent studies^32,50^, we were also able to demonstrate that the formation of 5’-dA (*i.e.*, 122.3 ± 13.1 µmol·min⁻¹·mg⁻¹ for **QCMT_1_** and 102.6 ± 4.5 µmol·min⁻¹·mg⁻¹ for **QCMT_2_**) and SAH (139.1 ± 36.7 µmol·min⁻¹·mg⁻¹ for **QCMT_1_** and 117.6 ± 4.2 µmol·min⁻¹·mg⁻¹ for **QCMT_2_**) paralleled the formation of **CH_3_-1Q**, consistent with the dual role of SAM as both a methyl donor and a radical initiator. As explained above, with the temperature used for the assay, 5’-dA and SAH quantifications were slightly underestimated due to their inherent instability in solution.

### QCMT alkylates the major groups of amino acid residues

Having developed a robust biochemical assay, we were able to investigate the substrate promiscuity of QCMT. A previous study with Mmp10 demonstrated strict substrate specificity for this B_12_-dependent radical SAM enzyme^26^. In contrast, **QCMT_1_** and **QCMT_2_** accepted all peptides with a modified functionalized residue, although their efficiencies varied based on the nature of the side chain (**Fig. 3a & b, Supplementary Figs. S4-S14**, and **Supplementary Table S2**). Notably, the closest homolog to **1Q**, the peptide **1N** (containing an Asn residue *in lieu* of Gln) proved to be an efficient substrate, along with peptides substituted with short hydrophobic residues such as Ala and Gly (**1A** & **1G**). More distantly related substrates with bulkier side chains (*e.g.*, Arg in **1R**), carboxyl groups (**1E**, **1D**), or aromatic groups (**1Y**) were surprisingly well accepted, particularly by **QCMT_2_**, with conversion yields >40%. MS/MS analysis confirmed that only the residue in position 7 was alkylated in each peptide assayed (**Supplementary Figs. S6-S14**). **QCMT_1_** exhibited a similar activity profile to **QCMT_2_**with negatively charged peptides, such as **1D**, being poorly methylated. Overall, these data show that while QCMTs display remarkable substrate promiscuity toward the functionalized residue, the nature of the side chain and the proximity of the functional group to the Cα atom affect catalytic efficiency.

**Figure 3.**
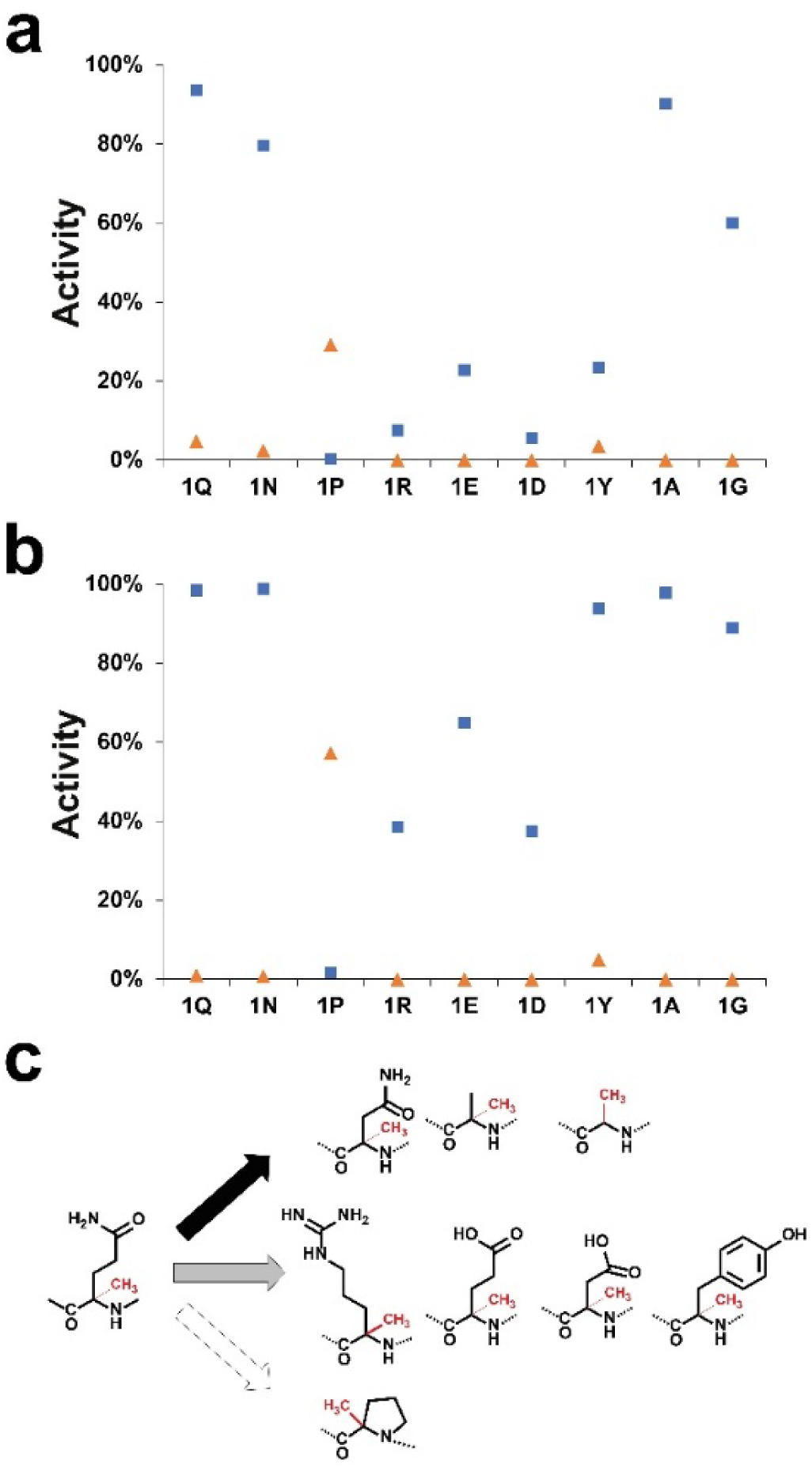
Activity of QCMTs on a library of synthetic peptides. (**a**) Activity of **QCMT_1_** and (**b**) **QCMT_2_** on 13mer peptides with various amino acid substitutions at the seventh position (Q, N, P, R, E, D, Y, A, and G). Percentages are given based on conversion into the methylated (blue squares) or epimerized (orange triangles) forms. **QCMT_1_** (7.5 µM) was incubated after iron-sulfur reconstitution with SAM (0.5 mM), peptide (150 µM), and OHCbl (0.1 mM) under anaerobic and reducing conditions for 30 minutes. Reactions were performed at 50°C and initiated by adding titanium citrate (2 mM). Single experiments. (*See* **Supplementary Figs. S4-S14 and Supplementary Tables S2-S4 for the full assignment**). (**c**) Structures of the different methylated amino acid residues formed by **QCMT_1_** and **QCMT_2_** during catalysis on 13-mer peptides.

Among all the peptides tested, the one containing a proline residue (**1P**) stood out. With this substrate, we obtained only trace amounts of methylated peptide ([M+3H]^3+^: 433.23), suggesting that the constrained environment of the Cα-atom hinders enzyme activity (**Fig. 3** and **Supplementary Figs. S4-S5**). However, this substrate was efficiently converted into another species with a molecular weight (**1P\***, [M+3H]^3+^: 428.56) identical to that of the substrate. This product was formed only in the presence of SAM, OHCbl, and a one-electron donor (Ti(III)), indicating that QCMT catalysed a substrate rearrangement. To determine the nature of this modification, the modified peptide (**1P\***) was purified, and its amino acid content was analyzed after acid hydrolysis and derivatization with *N*-α-(2,4-dinitro-5-fluorophenyl)-l-valinamide (L-FDVA) (**Fig. 4a**). The analysis was in agreement with its expected composition, except for the presence of large amount of D-Pro, as shown by its molecular weight ([M+H]^+^: 396.15) and retention time compared to an authentic standard (**Fig. 4a**, lower panel). Further confirmation of the composition of **1P\*** was obtained by comparison with an authentic peptide standard.

**Figure 4.**
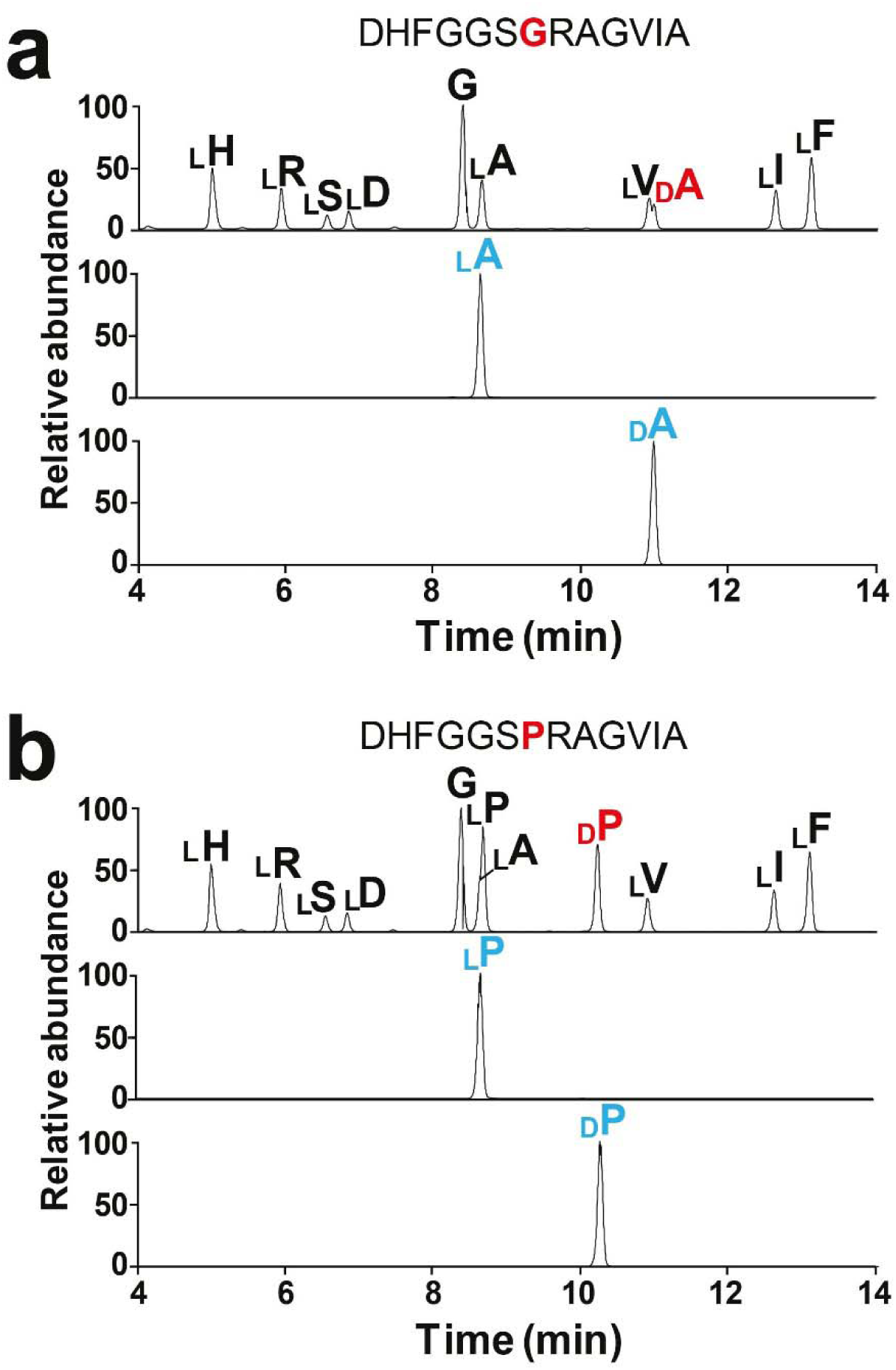
Analysis of the amino acid content of (a) 1P and (b) 1G after reaction with QCMT_1_. (upper traces: amino acid analysis, middle and lower traces: standards). Peptides were analyzed by acid hydrolysis, and derivatization with L-FDVA (see methods). Standards of derivatized L and D amino acids are indicated by (a) LA, DA, and (b) LP, DP (middle and lower panels). **QCMT_1_** (7.5 µM) was incubated after iron-sulfur reconstitution with SAM (0.5 mM), 1P or 1G (300 µM), and OHCbl (0.1 mM) under anaerobic and reducing conditions at 50°C in the presence of titanium citrate (2 mM).

A close analysis of the reactions revealed that small amounts of epimerized peptides were formed by both enzymes in the presence of **1N**, **1Y,** and **1Q**, mimicking the authentic substrate (**Fig. 3a** and **b**). These epimerized peptides were not further processed; notably, they were not alkylated, even when we performed fresh reactions with synthetic epimerized peptides. Interestingly, no epimerization occurred with **1A**, despite this peptide being efficiently methylated (**Fig. 3a**). Collectively, these results demonstrate that QCMT, in addition to its methylation activity, catalyzes the epimerization of amino acid residues similarly to radical SAM peptide epimerases: EpeE (T-SPASM group)^53^, PoyD and OpsD (One [4Fe-4S] cluster group)^54–57^. However, unlike these enzymes, this activity was strictly dependent on the presence of SAM, [4Fe-4S] cluster, and cobalamin. In addition, while radical SAM epimerases are active on short hydrophobic amino acid residues (*e.g.*, L-Leu, L-Ile, L-Val), QCMT has a unique substrate scope that includes amido- and aromatic amino acid residues.

### QCMT activity is independent of the presence of a side chain on the functionalized residue

Another surprising finding emerged from the analysis of the reaction with **1G** ([M+3H]^3+^: 415.21), in which Gln-7 was substituted by a Gly residue. This peptide proved to be an effective substrate for both **QCMT_1_**_&**2**_, as evidenced by its rapid conversion into a methylated peptide (**CH_3_-1G**, [M+3H]^3+^: 419.88) (**Fig. 3** and **Supplementary Figs. S4-S5**). LC-MS analysis of **CH_3_-1G** confirmed that a methyl group was transferred to the seventh residue, as expected (**Supplementary Fig. S14**). Further investigations revealed that Gly-7 was converted to an Ala residue with D-configuration (**Fig. 4b**).

The exclusive transformation of Gly into d-Ala underscores the strict control of stereochemistry exerted by QCMT, which catalyzes a reaction with a net retention of the configuration of the methylated carbon atom. This result also supports that QCMTs abstract the Gly pro-*R* hydrogen atom. Finally, similarly to peptides containing a D-Pro or D-Gln residue (*i.e.* **1P**, **1Q**), the d-Ala-containing peptide (**CH_3_-1G**) was not a substrate for QCMT, with notably no methylated product formed.

We then questioned whether the size of the substrate affects the reaction outcome (methylation *vs*. epimerization) and aimed to identify the minimal peptide size recognized by QCMT. A library of peptides was generated by truncating the *N*- and *C*-terminal ends. As shown, reducing the substrate size to as little as 7 residues had dramatic consequences on the reaction outcome (**Fig. 5**). Notably, while the peptide **2Q** remained an effective substrate, epimerization (**2Q*,** [M+2H]^2+^: 361.68) was favored over methylation (**CH_3_-2Q**, [M+2H]^2+^: 368.69) compared to **1Q** (**Fig. 5**). Here, the epimerized product (**2Q\***) accounted for approximately ∼45%, while **1Q\*** represented <5% (**Fig. 2b**) of the products formed. The nature of these products was confirmed through amino acid analysis and comparison with authentic standards. Like with larger substrates, QCMTs were unable to methylate a D-Gln (**Fig. 5**, **synth-2Q\***).

**Figure 5.**
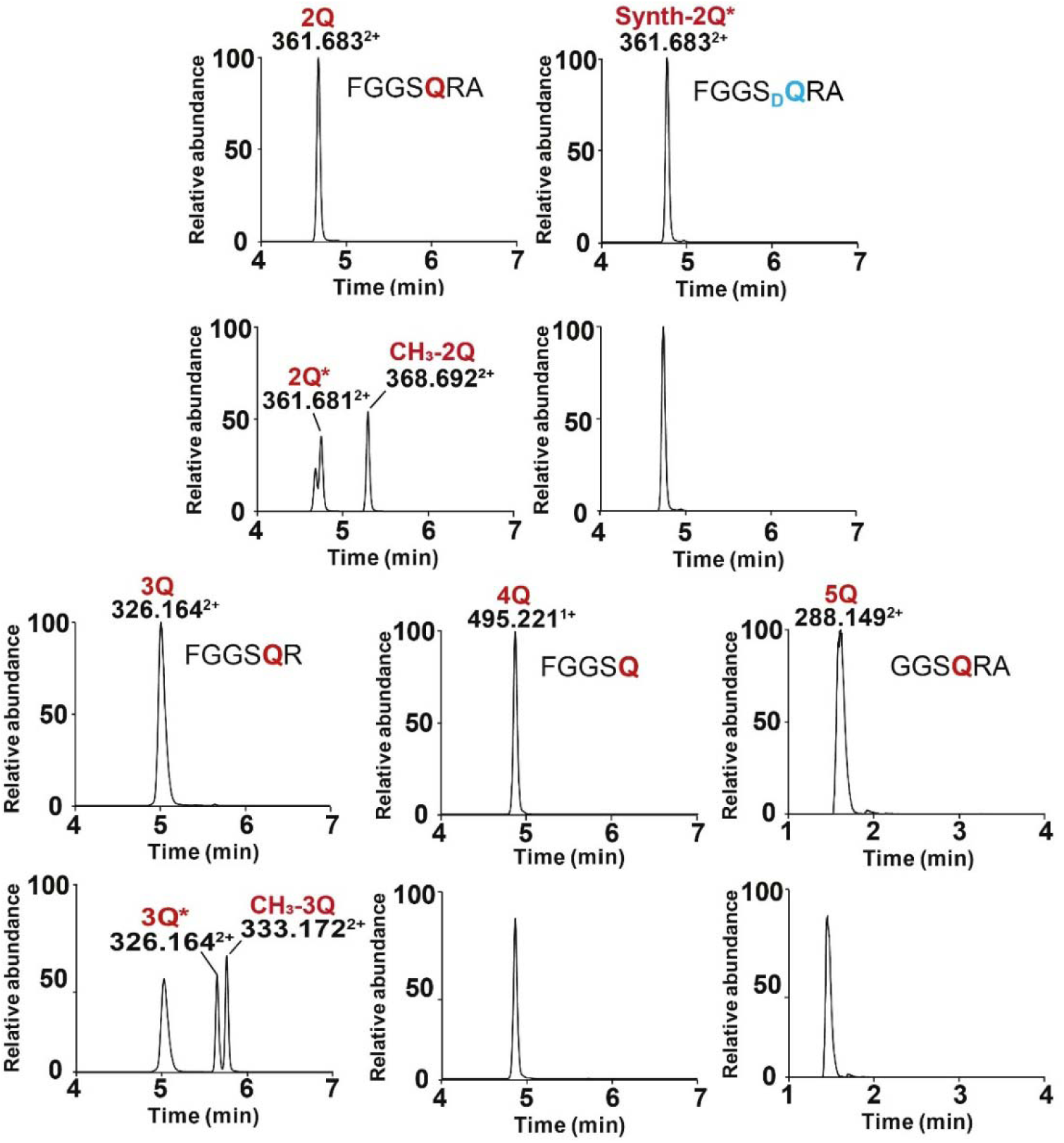
Influence of the peptide size (2Q to 5Q) on QCMT activity. **QCMT_1_** (7.5 µM) was incubated after iron-sulfur reconstitution with SAM (0.5 mM), peptide (150 µM), and OHCbl (0.1 mM) under anaerobic and reducing conditions. Reactions were performed at 50°C for 2 hours and initiated by adding T(III) (2 mM). See supplementary **figures S15-S17** and **Tables S2-S4** for the full assignment. **Synth-2Q\***: synthetic peptide containing a d-Gln.

Further truncation of the last *C*-terminal residue did not significantly change the reaction outcome with incubation of **QCMT_1_**with peptide **3Q**, resulting in the formation of an approximately 1:1 ratio of epimerized over methylated peptide. However, when the targeted glutamine residue was at the *C*-terminal end (*i.e.* **4Q**), it was no longer a substrate. Similarly, removing the *N*-terminal phenylalanine residue from **2Q** (resulting in **5Q,** [M+2H]^2+^: 288.15), resulted in a loss of activity. Collectively, these results demonstrate that QCMT requires only a few flanking residues for activity, establishing a minimalistic motif of six residues: FGGS**Q**R. Importantly, substituting the N-terminal phenylalanine with an alanine or a glycine residue did not restore enzyme activity (*data not shown*), indicating a critical role for this conserved residue, which interacts with the coenzyme F_430_ in MCR^58,59^.

### QCMT catalyzes C**α** H-atom abstraction and reversible H-atom abstraction

The epimerase activity of QCMT, although likely influenced by the *in vitro* conditions, is novel for a B_12_-dependent radical enzyme. Known radical SAM peptide epimerases typically require either a single catalytic [4Fe-4S] cluster ([4Fe-4S]_CAT_) or one [4Fe-4S]_CAT_ and one auxiliary cluster ([4Fe-4S]_AUX_) for catalysis^56,60,61^. These epimerases utilize a key cysteine residue to introduce a solvent-derived hydrogen atom, resulting in the formation of D-residues^53,54,61,62^. To investigate whether QCMT employs a similar mechanism, we first performed the reaction in a deuterated buffer. Under these conditions, the epimerized product **2Q^#^** ([M+2H]^2+^: 362.19) exhibited a +1 Da mass increment on the Gln residue, demonstrating that QCMT introduced a solvent-derived hydrogen atom in the epimerized product, like radical SAM epimerases (**Fig. 6a, 6b,** and **Supplementary Fig. S18**). A similar result was obtained with **1P**, leading to labeling of the proline residue (**Supplementary Fig. S19**). However, structural analysis, based on *in silico* models and the recently solved structure of MgmA^50^, does not support the use of an active-site cysteine residue to trap the radical intermediate. We therefore hypothesized that epimerization resulted from an uncontrolled reaction between the radical peptide intermediate and a buffer component. To validate this hypothesis, we performed the reaction under turnover-limiting conditions in a deuterated buffer. If epimerization is indeed the result of a free interaction between the radical intermediate and a solvent component, we would expect equal formation of deuterated D- and L-residues.

**Figure 6.**
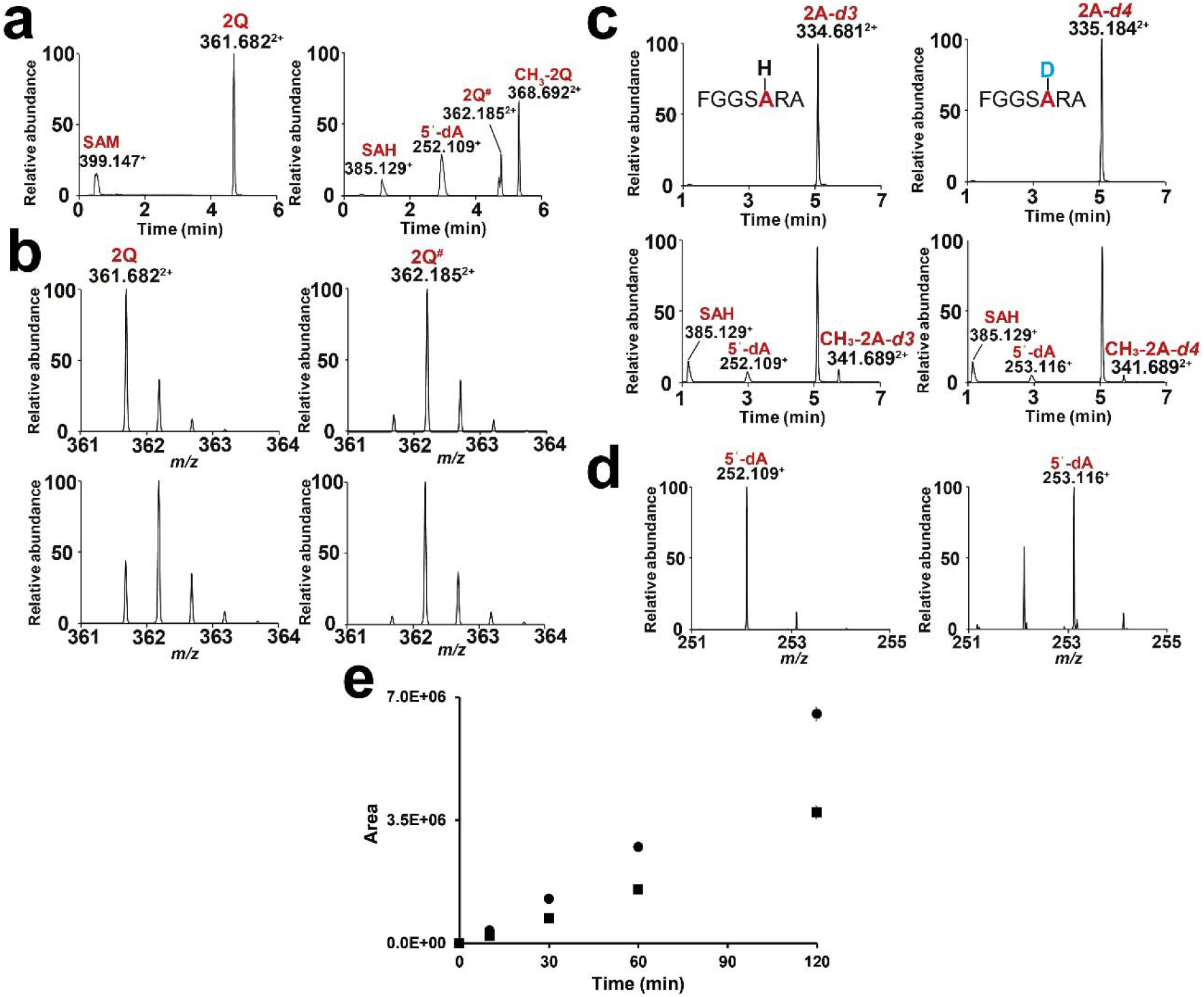
QCMT catalyzes reversible Cα H-atom abstraction. (a) LC-MS analysis of the reaction on **2Q** in deuterated buffer before (left panel) and after (right panel) 120 min incubation with **QCMT_1_**. (b) Isotopic distribution of peptides **2Q** (left panel) and **2Q^#^** (right panel) during incubation with **QCMT_1_** in deuterated buffer at T0 or 30 min, respectively (upper traces) and 120 min (lower traces). (c) LC-MS analysis of labeled peptides **2A-*d3*** (deuterated alanine on the methyl group) and **2A-*d4*** (per-deuterated residue) before (upper trace) and after (lower trace) incubation with **QCMT_1_**. (d) MS analysis of the 5’-dA production with peptide **2A-*d3*** (left panel) and **2A-*d4*** (right panel). (e) Time course analysis of the reaction catalyzed by **QCMT_1_** on peptides **2A-*d3*** and **2A-*d4***. Formation of the methylated peptides **2A-*d3*** and **2A-*d4*** is indicated by circle and square symbols, respectively. Experiments were performed in triplicate. **QCMT_1_** (7.5 µM) was incubated after iron-sulfur reconstitution with SAM (0.5 mM), peptide (150 µM), and OHCbl (0.1 mM) under anaerobic and reducing conditions. Reactions were performed at 50°C and started by adding Ti(III) (2 mM).

Under our assay conditions, the epimerized peptide **2Q^#^**was fully deuterated ([M+2H]^2+^: 362.19), while other products of the reaction, including 5’-dA and SAH, were not labeled. Strikingly, the [M+2H]^2+^: 362.19/361.68 ratio of **2Q** increased over time, with up to >66% deuterium incorporation at the end of the reaction (**Fig. 6b** & **Supplementary Figs. S20-S21**). Collectively, these findings support that the substrate radical intermediate reacted with a water-exchangeable buffer component, leading to uncontrolled hydrogen-atom addition. The radical intermediate, being likely planar, can react with equal probability on both faces leading to the formation of a labeled D- or L-residue.

To unequivocally demonstrate that QCMT catalyzes Cα hydrogen atom abstraction, we replaced the targeted Gln residue with a deuterated Ala residue, as has been done with other radical SAM enzymes^11,63,64^. Based on **2Q**, the shortest substrate for which an efficient activity was measured, we synthesized two novel peptides: one containing an Alanine with a deuterated side chain (**2A-*d3***) and the other with a per-deuterated residue (**2A-*d4***). Both peptides proved to be substrates; however, their conversion was significantly lower than that of **2Q** (**Fig. 6c)**. Incubation with **2A-*d4*** resulted in the formation of deuterated 5’-dA ([M+H]^+^: 253.11), whereas with **2A-*d3***, only unlabeled 5’-dA ([M+H]^+^: 252.11), was formed (**Fig. 6d**). These results unambiguously demonstrate that QCMT catalyzes Cα H-atom abstraction.

Taken together, our data revealed three unexpected findings. First, unlike Mmp10^26^, for which the side chain is critical for peptide interaction and modification, QCMT activity is only marginally affected by the nature of the side chain. Second, by incorporating Cα-methylation on a wide range of residues, QCMT enables the formation of unprecedented and unnatural post-translational modifications, including CαMe-Asn, CαMe-Arg, CαMe-Asp, CαMe-Glu, CαMe-Tyr, and CαMe-Ala. Finally, QCMTs, by converting a glycine residue into d-Alanine, catalyze a reaction currently not accessible by synthetic chemistry.

### Activity of QCMT with deuterated peptides 2A-*d3* and 2A-*d4*

As of now, the source of the hydrogen atom that is abstracted by B_12_-dependent radical SAM enzymes has been experimentally studied in five enzymes: GenK^48^, Fom3^46^, CysS^25^, TokK^47^, and Orf29^49^. In the case of TokK, and to a lesser degree with Orf29, a mixture of labeled and unlabeled 5’-deoxyadenosine (5’-dA) has been obtained. In contrast, GenK and Fom3 predominantly yielded mono-deuterated 5’-dA, with an observed mass of 253.11 Da, like QCMT (**Fig. 6d**).

For Orf29 and GenK, the isotopic distribution of the methylated products showed an unusual pattern characterized by significant deuterium enrichment. This phenomenon is thought to arise from a large kinetic isotope (KIE) and quantum tunneling effects^65^. Notably, GenK, the only B_12_-dependent radical SAM enzyme for which the influence of H-atom abstraction on the kinetic parameters has been investigated, displays a large KIE of 18 ±0.6. We took advantage of **2A-*d3*** and **2A-*d4*** to determine whether QCMT exhibited a similar KIE. As shown, with these labelled substrates, we measured only a modest KIE of approximately 1.8, indicating that hydrogen-atom abstraction is a partially rate-determining step (**Fig. 6e**). Furthermore, the methylated product displayed the expected isotopic distribution without any deuterium atom scrambling, in contrast to other radical SAM enzymes^8,48^.

## Discussion

Despite their discovery more than a decade ago, our understanding of B_12_-dependent radical SAM enzymes remains limited^1,21,66^, particularly regarding their unique ability to install methyl groups on unreactive *sp*^3^-hybridized carbon atoms, in a stereoselective manner. Recently, structures of several B_12_-dependent radical SAM enzymes have been solved^1^. Although these structures exhibit distinctive folds, they share notable similarities, particularly concerning the relative positions of the radical SAM [4Fe-4S] cluster and the cobalamin cofactor, which sandwich the enzyme’s substrates. Based on these structural insights, it has been proposed that B_12_-dependent radical SAM enzymes utilize a stepwise “pull-push” radical transfer mechanism, where 5’-dA “pulls” the hydrogen atom from one side of the substrate and MeCbl “pushes” a methyl radical from the opposite side^67^. In this mechanism, hydrogen-atom abstraction is predicted to be the rate-limiting step, while methyl transfer is exothermic.

We demonstrate here that QCMT uses SAM for both radical initiation and methyl transfer, as expected for a B_12_-dependent radical SAM methyltransferase catalyzing C*sp^3^*-alkylation. However, these two reactions can be dissociated, leading to hydrogen atom exchange and epimerization when the alkylation reaction is impaired. Unexpectedly, we found that QCMT exhibits exceptional substrate promiscuity, catalyzing post-translational modifications across all major classes of amino acid residues, including short hydrophobic residues, amido- and aromatic amino acid residues. This provides access to unnatural post-translational modifications, including the direct conversion of glycine residue into D-Ala, an unprecedented reaction. d-Ala is well-known in bacterial physiology and is typically formed by a two-step reaction involving serine dehydration and reduction^68^. To the best of our knowledge, QCMT is the only biocatalyst capable of converting Gly to d-Ala, making it a unique biochemical tool for engineering peptides and proteins by introducing unnatural post-translational modifications, for example, within short peptide tags. It also opens novel strategies for directly engineering MCR and for assessing the influence of novel post-translational modifications on their activity. Moreover, the hydrogen atom exchange and epimerase activity of QCMT is novel, opening the possibility of incorporating hydrogen atoms in late-stage synthesis and to epimerize aromatic and amido-amino acid residues.

Our study also reveals critical insights into the mechanism of B_12_-dependent radical SAM enzymes. We show that QCMT abstracts the Cα hydrogen atom, which is supported by labeling experiments and epimerization activity. The abstraction of a Cα hydrogen atom is a common theme across several classes of radical SAM enzymes, including glycyl radical activating enzymes (GRE-AE)^69,70^, thioether bond synthases^11,71^, and peptide epimerases^53,61^. All these enzymes are presumed to generate a Cα peptide radical intermediate, stabilized by the captodative effect. Although the presence of a side chain theoretically influences the planarity and stability of these Cα peptide radicals, recent studies indicate that peptide radicals can be unexpectedly stable^9^, consistent with QCMT’s hydrogen-atom exchange activity. Our study also demonstrates that QCMT only abstracts the pro-*R* H-atom, leading exclusively to the formation of d-Ala from Gly. This strict stereospecificity explains why QCMT, despite being largely insensitive to the amino acid side chain’s nature, is inactive toward d-amino acid residues (*e.g.,* d-Gln or d-Ala in **synth-2Q\*** and **CH_3_-1G**, respectively). Surprisingly, using deuterated substrates, we observed only a modest kinetic isotope effect, despite recent computational studies suggesting that hydrogen-atom abstraction should be the rate-limiting step^67^.

Our screening of QCMT homologues revealed that, even within the same enzyme family, some members, such as **QCMT_1_**, efficiently bind cobalamin, as shown by UV-visible and SEC-SAX analyses, whereas others, such as **QCMT_2_**, bind it reversibly. It remains to be clarified whether this observation is due to *in vitro* conditions or if it has physiological relevance. Indeed, we showed that in solution, cobalamin binding influences the overall shape of the protein, resulting in a more compact and closed conformation.

At the mechanistic level, our data indicate that hydrogen-atom abstraction and methyl-transfer reactions are not concerted. We propose that the radical peptide intermediate reorients during catalysis to attack the methyl group in a stereo-controlled, but not concerted, manner. The observation that the amino acid side chain plays a minor role in the reaction rate suggests that there are limited constraints within the active site. This contrasts with Mmp10, where a dense network of interactions stabilizes the substrate’s side chain within the active site^26^.

In conclusion, we have demonstrated that while QCMT catalyzes a strictly controlled stereo- and regio-selective reaction, it also exhibits unexpected catalytic promiscuity, leading to peptide epimerization, hydrogen atom exchange, and alkylation (**Fig. 7**). QCMT is a promising radical biocatalyst for accessing novel peptide structures with unexpected reactivity.

**Figure 7.**
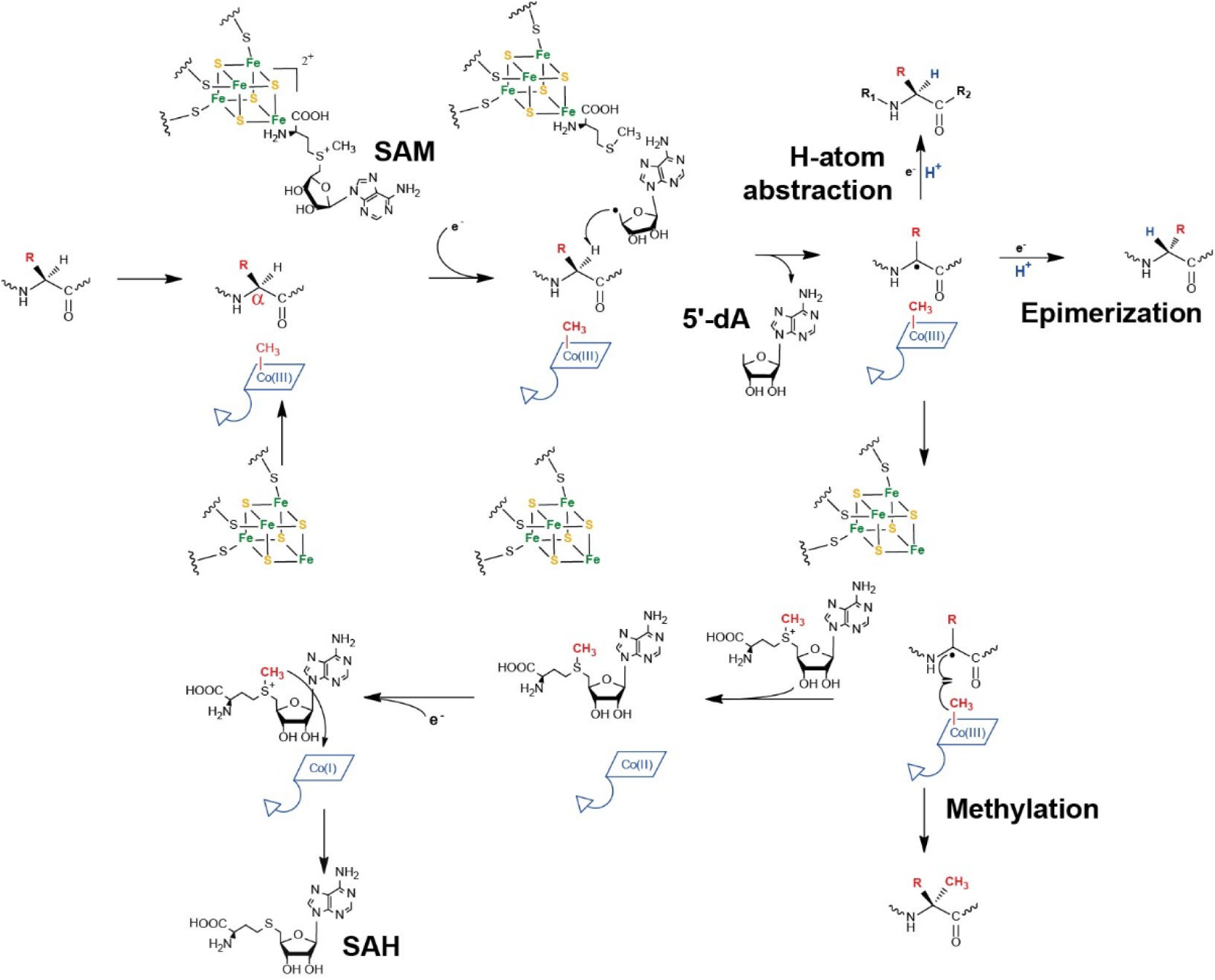
Proposed mechanisms for QCMT leading to methylation, epimerization or reversible H-atom abstraction. QCMT catalytic promiscuity: Following Cα H-atom abstraction, a carbon-centered intermediate is formed that either reacts with MeCbl or a solvent-derived H-atom, leading to methylation, labelling (reversible H-atom abstraction), and epimerization reaction. In the absence of a side-chain (**R**), methylation leads to the conversion of Gly into d-Ala.

## Authors’ Contributions

The manuscript was written through the contributions of all authors. All authors have given approval to the final version of the manuscript.

## Funding Sources

This work was supported by ANR (grants ANR-20-CE44-0005, ANR-21-CE11-0030, ANR-23-CE07-0046).

## Acknowledgments

We acknowledge SOLEIL (Saint-Aubin, France) for the provision of synchrotron radiation facilities (proposal 20221418), and we would like to thank the SWING staff for assistance in using the beamline.

## Data availability

All data generated or analyzed during this study are included in the text and figures of this article and its Supplementary Materials. All other data are available from the corresponding author upon reasonable request.

## Ethics declarations

### Competing interests

The authors declare no competing interests.

## Notes

### Competing Interest Statement

The authors have declared no competing interest.

