## Supplementary Information for "The stereochemical mechanism of the B_12_-dependent radical SAM glutamine methyltransferase (QCMT): Novel insights and unprecedented post-translational modifications"

c. Aix Marseille Univ, CNRS, IM2B, IMM, BIP UMR 7281, Marseille, France

d. Synchrotron SOLEIL, HelioBio group, L'Orme des Merisiers, Départementale 128, 91190 Saint-Aubin, France

Figure S9 - MS-MS fragmentation spectra of peptides 1R (upper panel) and $CH_3$-1R (lower panel). .. 12
Figure S10 - MS-MS fragmentation spectra of peptides 1E (upper panel) and $CH_3$-1E (lower panel).. 13
Figure S11 - MS-MS fragmentation spectra of peptides 1D (upper panel) and $CH_3$-1D (lower panel). 13
Figure S12 - MS-MS fragmentation spectra of peptides 1Y (upper panel) and $CH_3$-1Y (lower panel).. 13
Figure S13 - MS-MS fragmentation spectra of peptides 1A (upper panel) and $CH_3$-1A (lower panel). 14
Figure S14 - MS-MS fragmentation spectra of peptides 1G (upper panel) and $CH_3$-1G (lower panel). 14

### Material and methods

#### Cloning of QCMT homologs

The genes of QCMT from: *Methanoculleus thermophilus* ($QCMT_1$): WP_066957658, *Methanothermococcus thermolithotrophicus* ($QCMT_2$): WP_018153439, *Methanoculleus marisnigri*: A0A117MFC8, *Methanothermococcus okinawensis*: WP_013866670 and *Methanoculleus bourgensis*: WP_074174763, were produced by Genecust with codon optimization for *Escherichia coli* expression. Genes were cloned into pET28a(+) plasmid with a *N*-terminal Strep-tag sequence. Plasmids were then transformed into *E. coli* BL21 Star (DE3) cells (Life Technologies) for protein expression.

#### Expression and protein purification

*E. coli* cells transformed with the corresponding plasmids were grown in LB medium supplemented with Ampicillin (0.1 ug.mL$^{-1}$). Cultures were incubated at 37 °C until $OD_{600}$ reached 0.8 (~4 h, 160 rpm), then 0.3 mM of ammonium iron(II) citrate was added and protein expression was induced by adding 0.5 mM isopropyl-β-d-1-thiogalactopyranoside (IPTG). The culture was further incubated (16 h, 21 °C, 160 rpm) before cells are harvested by centrifugation (10 min, 4 °C, 5500 *x* g). The cell pellet was washed with cold buffer (250 mL, 50 mM Tris, 300 mM NaCl, 10% glycerol pH 8) then frozen in liquid nitrogen and stored at -80 °C until further use.

Pellet cell was resuspended (300 mL for 9 L of culture volume) in lysis buffer (50 mM Tris, 300 mM NaCl, 10% glycerol pH 8) containing complete ULTRA EDTA-free protease inhibitor tablets (Roche), 0.5% Triton X-100, and 10 mM β-mercaptoethanol. Lysozyme was then added and the suspension (1 mg.mL$^{-1}$) was incubated for 30 min at room temperature before cells were lysed by sonication at 60% amplitude (8 × 45 s with 60 s cooling intervals on ice). The lysate was ultracentrifuged (45 min, 4 °C, 150 000 × g), and the supernatant was kept on ice. All subsequent steps were carried out at 4 °C. Purification of the crude extract, containing the recombinant protein **$QCMT_1$** or **$QCMT_2$** was purified by affinity chromatography on Strep-Tactin® Superflow® high-capacity resin (IBA-Lifesciences) at a flow rate of 1 mL min$^{-1}$ equilibrated with lysis buffer. The crude extract was applied to the Strep-tag® column and the resin was then washed with 100 mL of lysis buffer. The Strep-tagged protein was eluted (1 mL.min$^{-1}$) with *D*-desthiobiotin (3 mM) and dithiothreitol (3 mM) in lysis buffer. Fractions containing the protein are pooled and concentrated on a 30 kDa cutoff filter and subsequently frozen in liquid nitrogen and stored at -80 °C for further use. The purity of the **$QCMT_1$** or **$QCMT_2$** proteins was verified by SDS-PAGE electrophoresis. Concentrations of protein were estimated by using spectrophotometer (**$QCMT_1$**: $M_w$ = 60868.31 Da, ε = 87320 M$^{-1}$ cm$^{-1}$; **$QCMT_2$**: $M_w$ = 60804.19 Da, ε = 87320 M$^{-1}$ cm$^{-1}$).

#### Reconstitution of QCMTs

All protein reconstitutions and experiments were performed under anaerobic conditions, in a glovebox. Reconstitutions were performed by adding 3 mM DTT, followed by a six-fold molar excess of $(NH_4)_2Fe(SO_4)_2$ and $Na_2S$. Precipitated iron sulfides were removed by using a desalting column (PD 10 Desalting Columns, Cytiva) equilibrated with buffer (50 mM Tris, 300 mM NaCl, 10% glycerol pH 8). The protein was concentrated by ultrafiltration (30 kDa MWCO).

#### Peptide synthesis

Peptides **1Q** to **5Q** (*see* Table S2) were synthetized by Proteogenix. Peptides **1A-*d3*** and **1A-*d4*** were synthetized by PRIMACEN platform, Université de Rouen Normandie France. Long peptides (**1Q** to **1G**, Table S2) were solubilized in a 0.05% Formic acid solution. Short peptides (**2Q** to **1A-*d4***, Table S2) were solubilized in a 50 mM Tris, 300 mM NaCl pH 8 buffer. The pH of each short peptide solution was measured and adjusted to 7-8.

Peptide synthesis of **1A-*d3*** and **1A-*d4*** were performed by Fmoc solid phase methodology on a Liberty microwave assisted automated peptide synthesizer (CEM) using the standard manufacturer procedures at 0.1 mmol scale on preloaded Fmoc-Ala-Wang resin. The synthetic peptides were deprotected, cleaved from the resin with a mixture of TFA/TIS/$H_2O$ (9.5:0.25:0.25) and precipitated in *tert*-butylmethyl ether, and then purified by reversed-phase (RP) HPLC (Gilson) on a 22 x 250 mm Vydac 218T1022 $C_{18}$ (10 µm, 300 Å) column (Grace) using a linear gradient (5-30% over 45 min) of acetonitrile/TFA (99.9:0.1) at 10 mL.min$^{-1}$. Characterization of the purified peptides was then achieved by MALDI-TOF mass spectrometry on a UltrafleXtreme (Bruker) in the reflector mode using α-cyano-4-hydroxycinnamic acid as a matrix. Analytical RP-HPLC, using a 4.6 x 250 mm Vydac 218T54 $C_{18}$ (5 µm, 300 Å) column, indicated that a peptide purity > 99.9%.

**Enzymatic assay**

All assays were performed under anaerobic conditions in a glovebox and in the dark, at 50°C. Activity assays were done using freshly prepared reconstituted protein (7.5 µM). All reagents were solubilized in water in the glovebox just before using them. Protein was incubated first with 100 µM of OH-cobalamin at 4°C, then mixed successively with 100 mM Tris pH8, 1 mM DTT, 500 µM SAM (Sigma-Aldrich), 150 µM peptide, and 2 mM titanium(III) citrate (final concentrations). A sample was collected at T = 0 min before the addition of titanium(III) citrate, and a second at T = 30 min (long peptides) or T = 120 min (short peptides).

For enzyme kinetics, the concentration of SAM and peptide was increased to 800 µM and 150 µM, respectively, in order to measure linear activity over a longer period of time. Samples were collected at 5 min, 10 min, 20 min, 30 min and 60 min. All samples were quenched by a 10-fold dilution in 0.3% TFA.

**LC–MS analysis**

Peptides modification characterization was done by liquid chromatography on a Vanquish UHPLC systems coupled to a Q Exactive Focus mass spectrometer (Thermo Fisher Scientific). Quenched samples, after 15 times dilution in formic acid 0.1% (buffer A), were injected (2 µL) on a reversed-phase Phenyl-Hexyl column (2.1 mm x 50 mm, 1.8µM, Agilent). Optimization of peptide analysis was performed in order to have the best separation possible, especially between peptides and epimerized peptides. All compounds were eluted at a flow rate of 0.3 mL.min$^{-1}$ with a gradient from 0% to 18% (large peptides) or from 0% to 12% (short peptides) of buffer B (acetonitrile 80% and formic acid 0.1%).

To characterize the post-translational modifications of each peptide, fragmentation was achieved on the most abundant ion (Tables S2 and S3). Collision energie of 22% was used for peptide **1Q** to **1G,** 20% for peptides **2Q** to **1A-d4** and 15% for peptides **3Q** to **5Q**. LC-MS spectras and Isotopic distributions were generated using the Qual Browser software (Thermo Xcalibur, v2.0.3). All compounds were analyzed by mass spectrometry in positive mod with a mass range between 200 and 1000 uma. LC-MS/MS spectra were manually deconvoluted and annotated using the FreeStyle software with the Xtract tool (ThermoFisher, v1.3). When signal intensity was too low for the deconvolution, Qual Browser software was used to generate the spectra (**Figs. S15** and **S17**).

**HPLC analysis**

Quantification of compounds from *in vitro* assays was performed with an Agilent Technologies 1200 series HPLC using calibration curves. Aliquots (10 µL) of standards and quenched samples were injected onto a LiChrospher® 100 RP-18 endcapped column (250 x 4 mm). Elution was performed using an acetonitrile gradient (18.7–60% solvent B) at 1 mL.min$^{-1}$. Solvent A was 0.1% TFA; solvent B was 80% acetonitrile + 0.1% TFA. UV detection was at 215 nm for peptides and 260 nm for SAH/5'-dA.

**Peptide purification and L-FDVA derivatization**

To determine the amino acid composition of peptides **1P**, **1G** and **2Q**, large-scale reactions (500µL) were performed. Enzymes were removed by adding 50 µL of 10% TFA, then diluting with 450 µL of 0.1% TFA. After 10 min incubation on ice (between each addition of TFA) and centrifugation, peptides were purified on SepPak® C18 cartridges and eluted with 80% acetonitrile + 0.1% TFA.

Eluates were dried in borosilicate glass tubes (Kimble, ASTM type 1) for 1 h in a Savant SPD121P SpeedVac. Hydrolysis was performed by adding 1 mL of 6 M HCl, applying partial vacuum (<1 mbar), and incubating at 150 °C for 1 h. Hydrolysates were derivatized with 10 µL $NaHCO_3$ 1 M, 10 µL L-FDVA (10 µg µL$^{-1}$ in acetonitrile), 10 µL acetonitrile, and 10 µL water, incubated at 42 °C for 1 h. Derivatized samples were analyzed by LC–MS using a Hypersil GOLD aQ column (2.1 × 200 mm, 1.9 µm). Compounds were eluted between 30–70% buffer B at 0.3 mL min$^{-1}$. Glutamine is converted to glutamic acid during acid hydrolysis; therefore, L- and D-glutamic acid contents were assessed using synthetic peptide Synth-2Q containing D-glutamine.

**UV-visible spectroscopy**

The UV-Visible spectra were recorded with a protein concentration of 1 mg mL$^{-1}$ in a quartz cuvette with an optical path length of 1 cm under anaerobic conditions on a Jasco V-760 spectrophotometer using a scan rate of 40 nm min$^{-1}$. UV–visible spectra (215–700 nm) were recorded before and after protein reconstitution and incubation with 200 µM OH-cobalamin. Excess OH-cobalamin was removed using a PD-10 column (Cytiva).

**SEC-SAXS analysis**

Reconstituted and tag-cleaved **QCMT$_1$** was purified by size exclusion on a Superdex 200 Increase 10/300 GL column (GE Healthcare) using 50 mM Tris, 300 mM NaCl, 10% glycerol pH 8 buffer containing 1 mM DTT to prevent aggregation. Protein samples (≈100 µM and 20 µM respectively) were incubated with or without 200 µM OHCbl and analyzed by SEC–SAXS using an in-line Superdex 200 Increase 5/150 GL column (GE Healthcare). Samples were loaded anaerobically onto an HPLC system equilibrated with degassed buffer (50 mM Tris, 300 mM NaCl, 5% glycerol pH 8). SAXS data were collected on the SWING beamline at Synchrotron SOLEIL (Saint-Aubin, France). Acquisition and data-processing parameters are summarized in **Supplementary Table S5.**

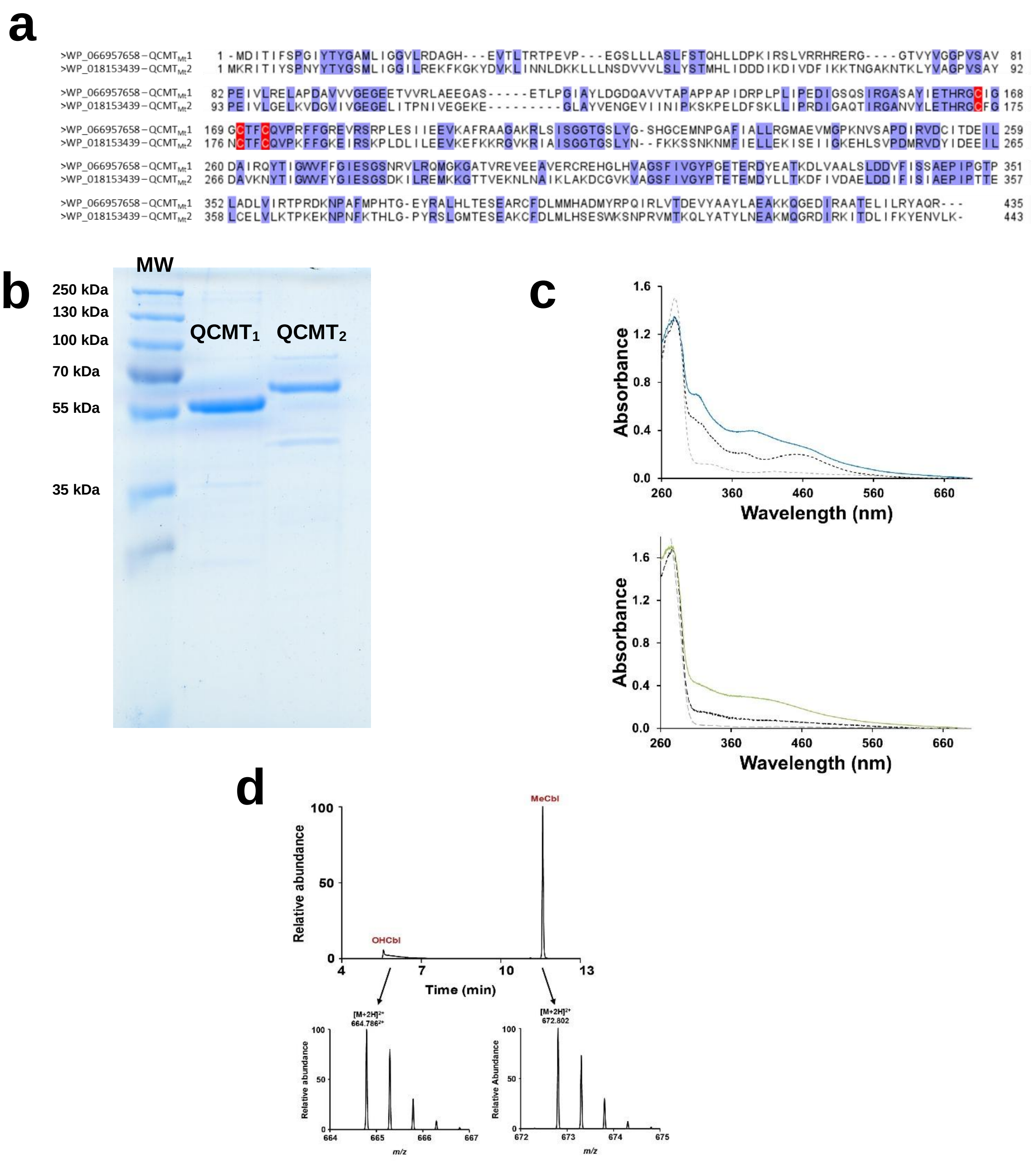

**Figure S1 –** Sequences (**a**), SDS-PAGE (b) and UV-visible analysis of **$QCMT_1$** and **$QCMT_2$**.

(**a**) Sequence alignment of **$QCMT_1$** and **$QCMT_2$**. Strictly conserved amino acid residues are highlighted in blue, and conserved cysteine residues of the radical SAM motif are highlighted in red. (**b**) SDS-PAGE analysis of **$QCMT_1$** and **$QCMT_2$**. MW: molecular weight markers (*Uncropped gel see Supplementary Figure S22*). (**c**) UV-vis spectra of QCMT from *Methanoculleus thermophilus* (**$QCMT_1$**: upper panel) and from *Methanothermococcus thermolithotrophicus* (**$QCMT_2$**: lower panel). Dotted grey line: as purified enzyme; dotted black line: enzyme after incubation with MeCbl and gel filtration; solid line: enzyme after incubation with MeCbl, gel filtration and iron-sulfur reconstitution. (**d**) LC-MS analysis of the cobalamin content of **$QCMT_1$** after reconstitution with MeCbl. As shown **$QCMT_1$** contains MeCbl ($[M+2H]^{2+}$: 672.08 and OHCbl ($[M+2H]^{2+}$: 664.78).

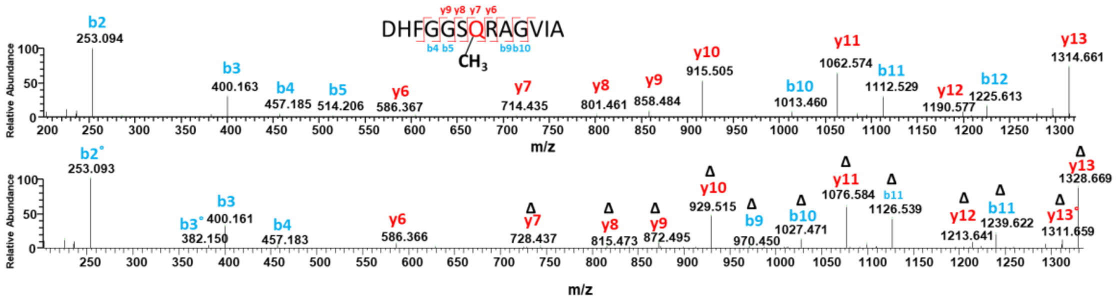

**Figure S2 -** MS-MS fragmentation spectra of peptides **1Q** (upper panel) and **$CH_3$-1Q** (lower panel).

Ions with a loss of $H_2O$ are noted by ˚.

Ions with an addition mass of +14.01565 Da corresponding to an addition of a $CH_2$ are noted by $^{\Delta}$.

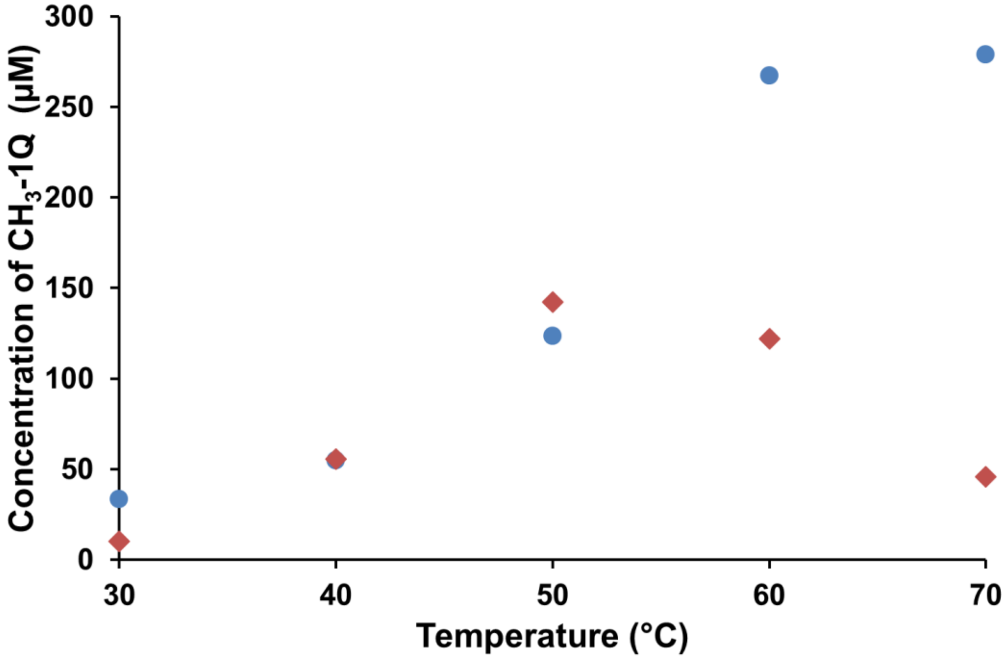

**Figure S3 –** Activity of **QCMT$_1$** (blue) and **QCMT$_2$** (orange) depending on incubation temperature.

Activity is based on the formation of methylated peptide ($CH_3$-1Q) after 20 min incubation. Incubation was performed under anaerobic conditions in the presence of enzyme (7.5 µM), SAM (0.8 mM), OHCbl (100 µM), titanium (III) citrate (2 mM) and 1Q (300 µM). Single experiments.

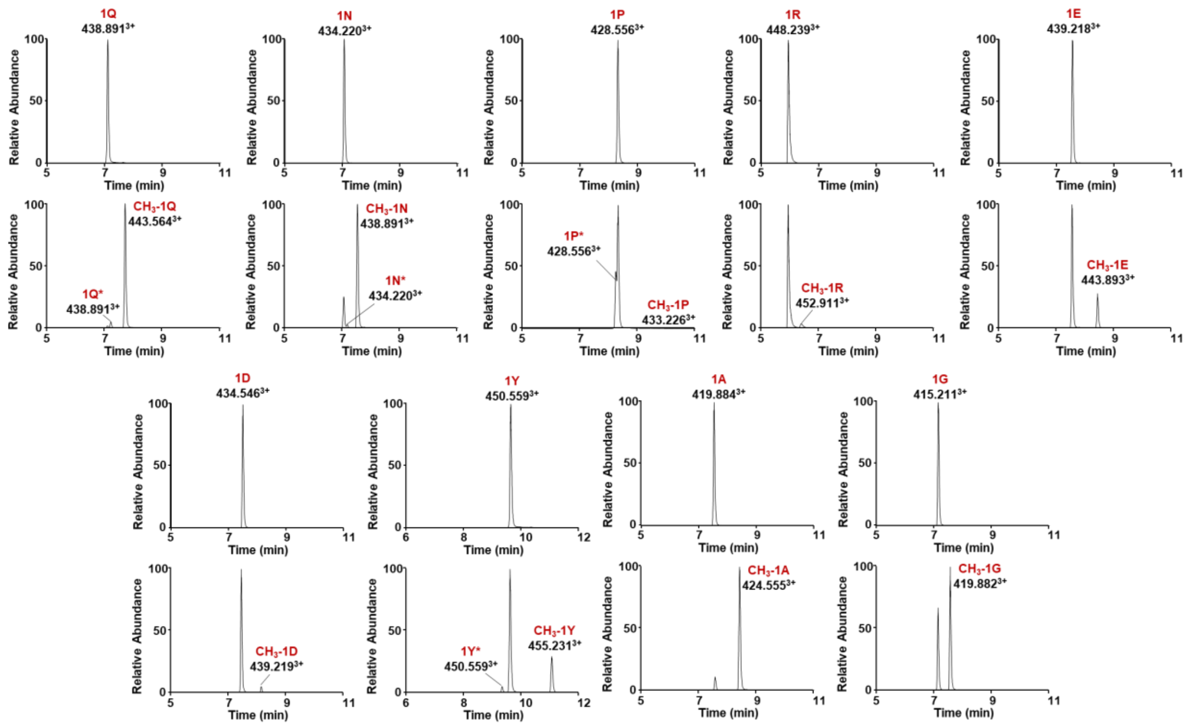

**Figure S4 –** Activity of **QCMT$_1$** on peptides **1Q**, **1N**, **1P**, **1R**, **1E**, **1D**, **1Y**, **1A**, **1G.**

QCMT$_1$ (7.5 µM) was incubated after iron-sulfur reconstitution with 150 µM peptides, 0.5 mM SAM and 0.1 mM OHCbl under anaerobic and reducing conditions. Reactions were performed at 50°C, initiated by adding titanium citrate (2 mM) and incubated for 30 minutes. * indicates epimerized peptide.

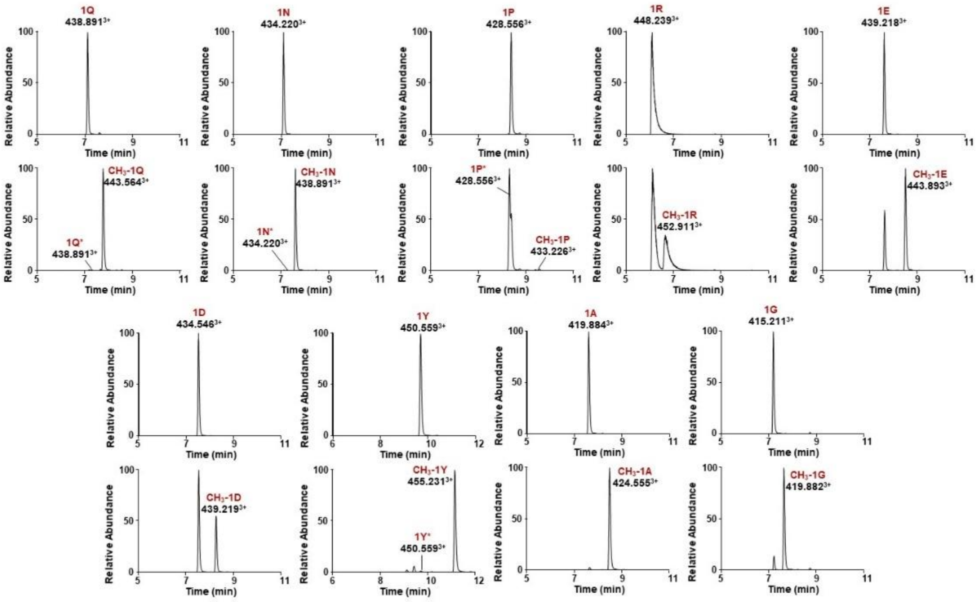

**Figure S5 –** Activity of **QCMT$_2$** on peptides **1Q**, **1N**, **1P**, **1R**, **1E**, **1D**, **1Y**, **1A**, **1G.**

**QCMT$_1$** (7.5 µM) was incubated after iron-sulfur reconstitution with 150 µM peptides, 0.5 mM SAM and 0.1 mM OHCbl under anaerobic and reducing conditions. Reactions were performed at 50°C initiated by adding titanium citrate (2 mM) and incubated 30 minutes.

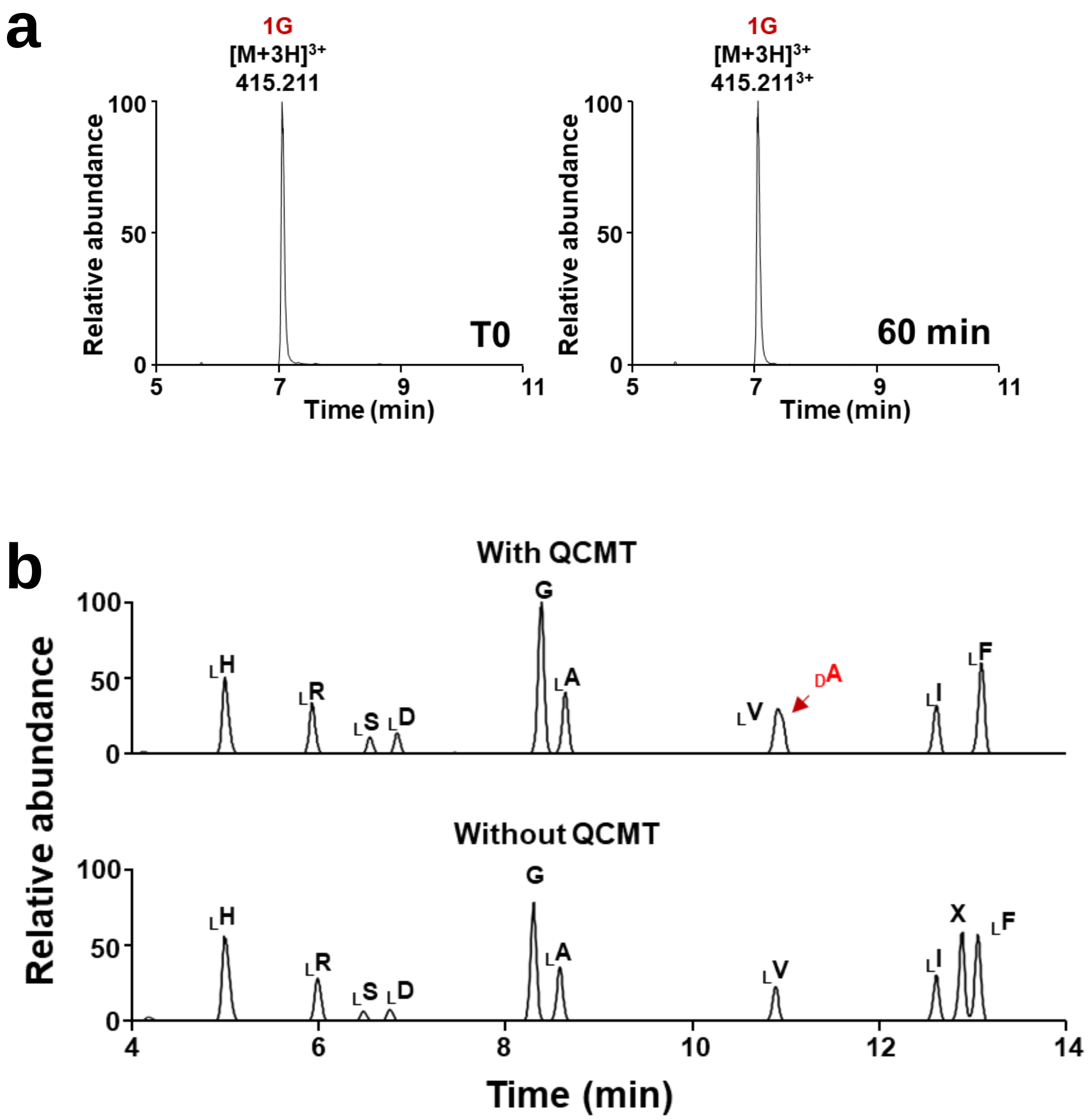

**Figure S6 – Control experiment with 1G.**

(**a**) LC-MS analysis of peptide **1G** (300 µM) incubated in the presence of SAM (0.8 mM), MeCbl (0.1 mM) and titanium citrate (2 mM), under anaerobic and reducing conditions at 50°C. (**b**) Analysis of the amino acid content of **1G** after reaction with **$QCMT_1$** (upper traces) or in the absence of **$QCMT_1$** (lower trace). Amino acid content was determined by LC-MS analysis, after acid hydrolysis of the corresponding peptides and derivatization with L-FDVA. X denotes an impurity.

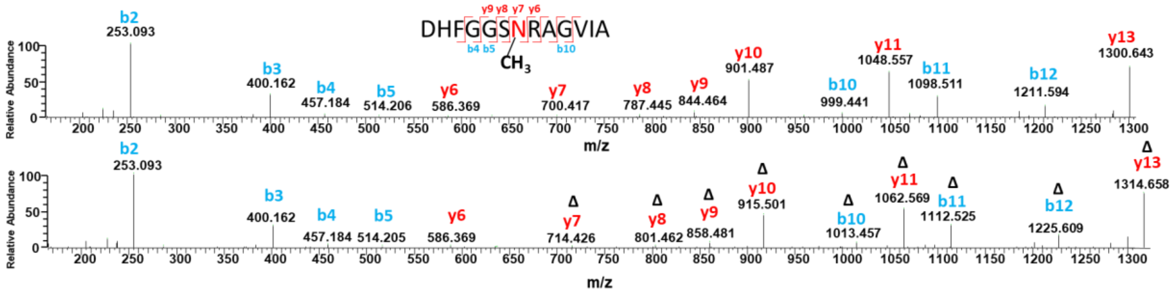

**Figure S7 -** MS-MS fragmentation spectra of peptides **1N** (upper panel) and **$CH_3$-1N** (lower panel).

Ions with a mass addition of +14.01565 Da are noted by $^{\Delta}$.

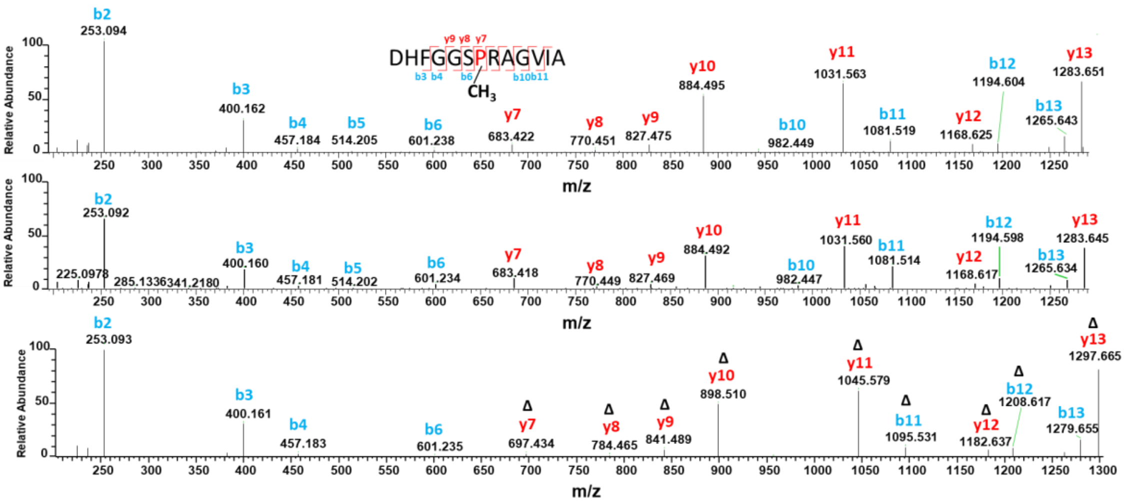

**Figure S8 -** MS-MS fragmentation spectra of peptides **1P** (upper panel), **1P*** (middle panel) and **$CH_3$-1N** (lower panel).

Ions with a mass addition of +14.01565 Da are noted by $^{\Delta}$.

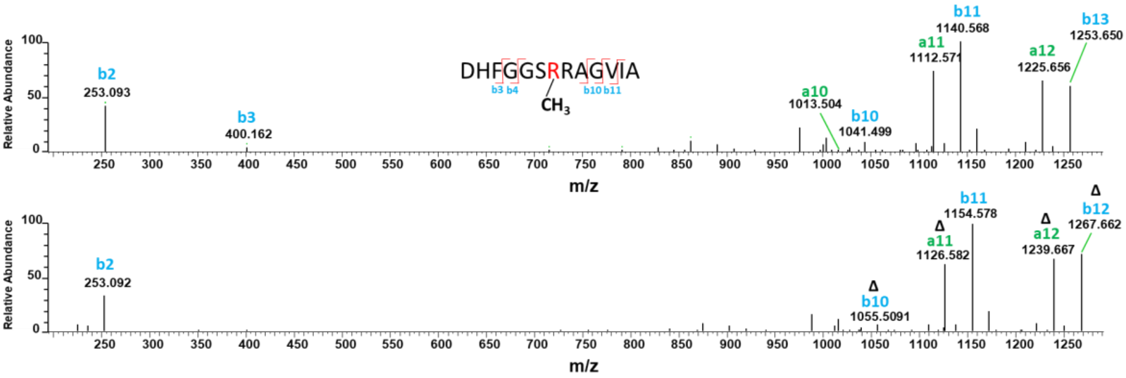

**Figure S9 -** MS-MS fragmentation spectra of peptides **1R** (upper panel) and **$CH_3$-1R** (lower panel).

Ions with a mass addition of +14.01565 Da are noted by $^{\Delta}$.

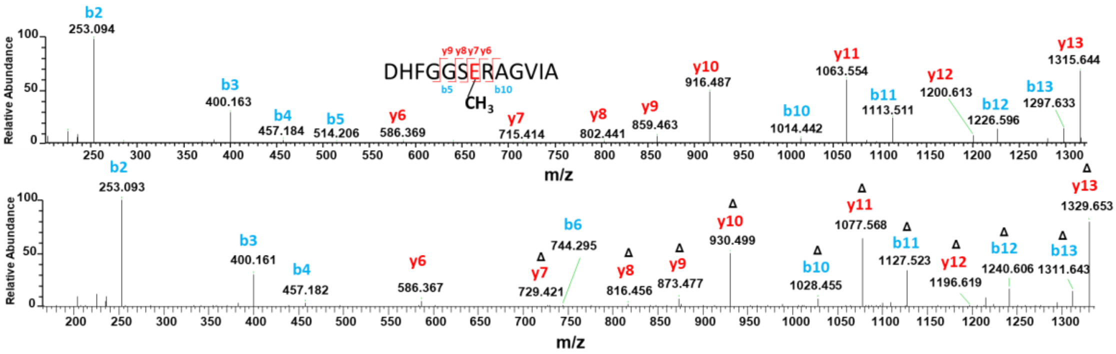

**Figure S10 -** MS-MS fragmentation spectra of peptides **1E** (upper panel) and **$CH_3$-1E** (lower panel). Ions with a mass addition of +14.01565 Da are noted by Δ.

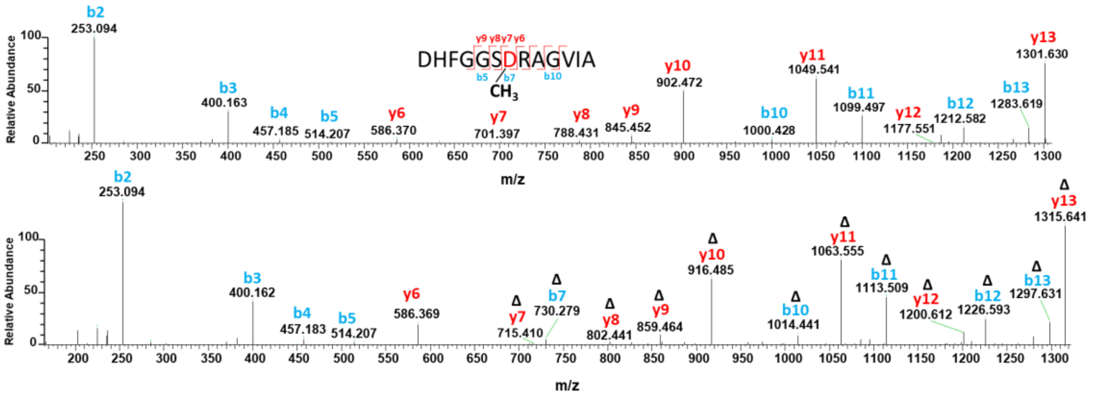

**Figure S11 -** MS-MS fragmentation spectra of peptides **1D** (upper panel) and **$CH_3$-1D** (lower panel). Ions with a mass addition of +14.01565 Da are noted by Δ.

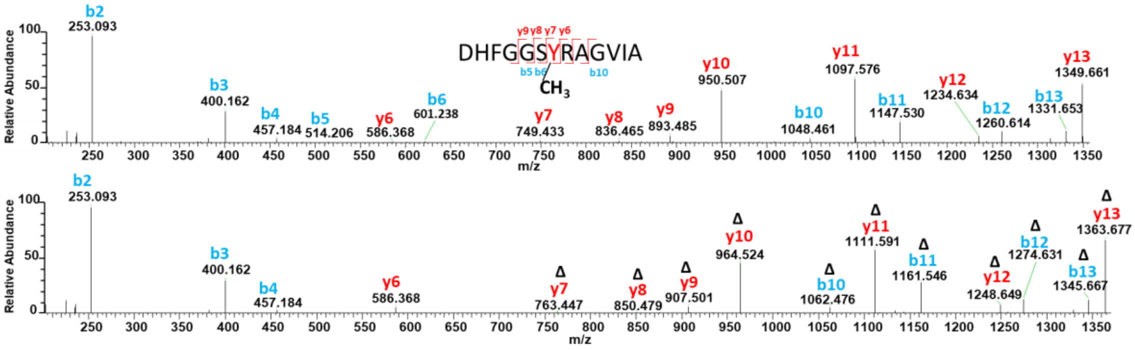

**Figure S12 -** MS-MS fragmentation spectra of peptides **1Y** (upper panel) and **$CH_3$-1Y** (lower panel). Ions with a mass addition of +14.01565 Da are noted by Δ.

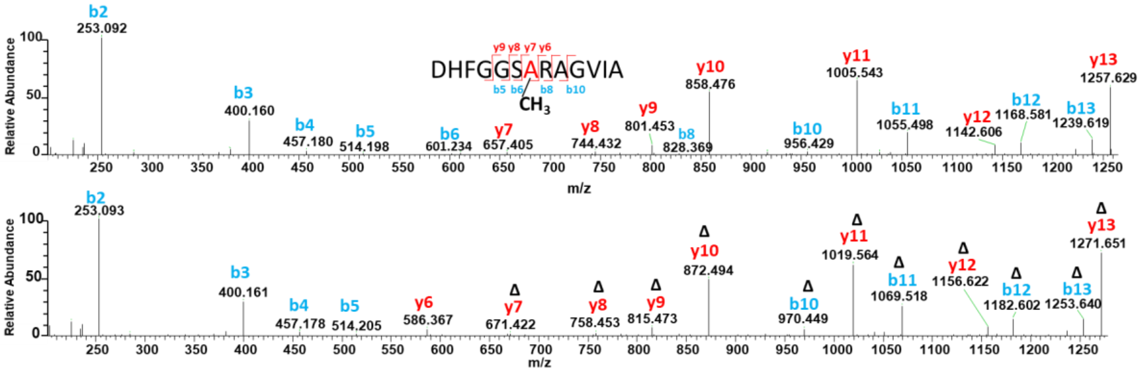

**Figure S13 -** MS-MS fragmentation spectra of peptides **1A** (upper panel) and **$CH_3$-1A** (lower panel). Ions with a mass addition of +14.01565 Da are noted by Δ.

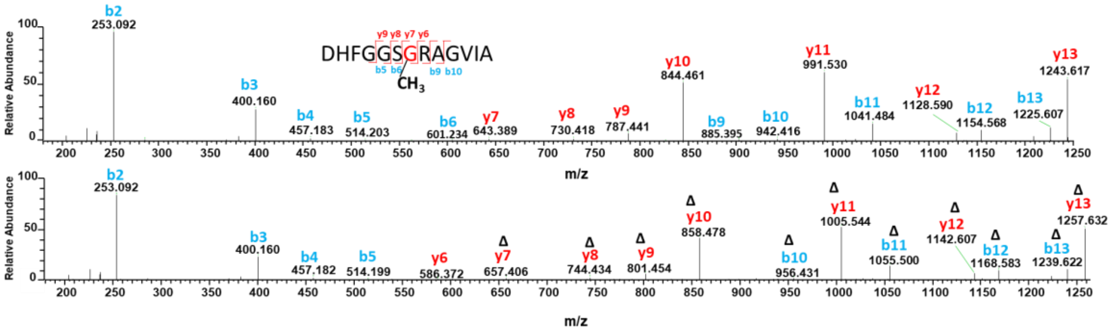

**Figure S14 -** MS-MS fragmentation spectra of peptides **1G** (upper panel) and **$CH_3$-1G** (lower panel). Ions with a mass addition of +14.01565 Da are noted by Δ.

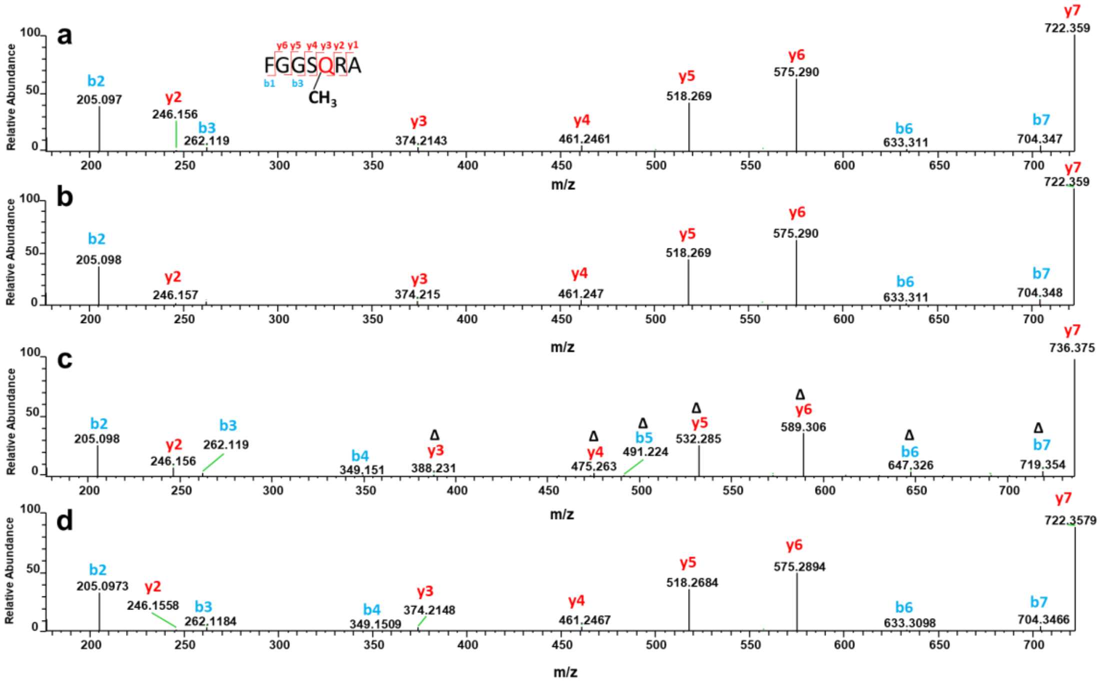

**Figure S15** - MS-MS fragmentation spectra of **2Q** (panel a), **2Q*** (panel b), **$CH_3$- 2Q** (panel c) and **Synth-2Q*** (panel d).

Ions with a mass addition of +14.01565 Da are noted by Δ.

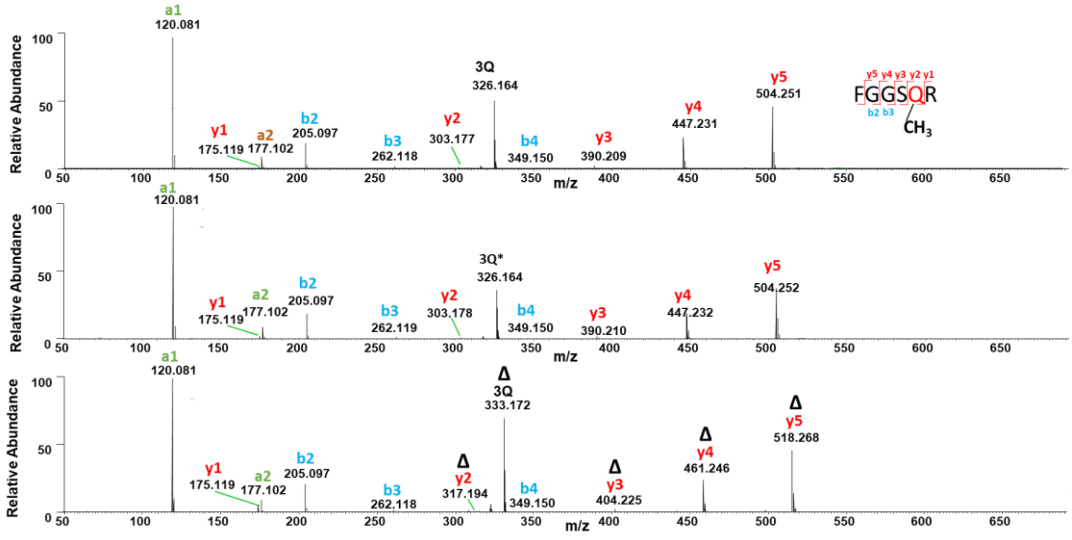

**Figure S16** - MS-MS fragmentation spectra of **3Q** (upper panel), **3Q*** (middle panel) and **$CH_3$-3Q** (lower panel).

Ions with a mass addition of +14.01565 Da are noted by Δ.

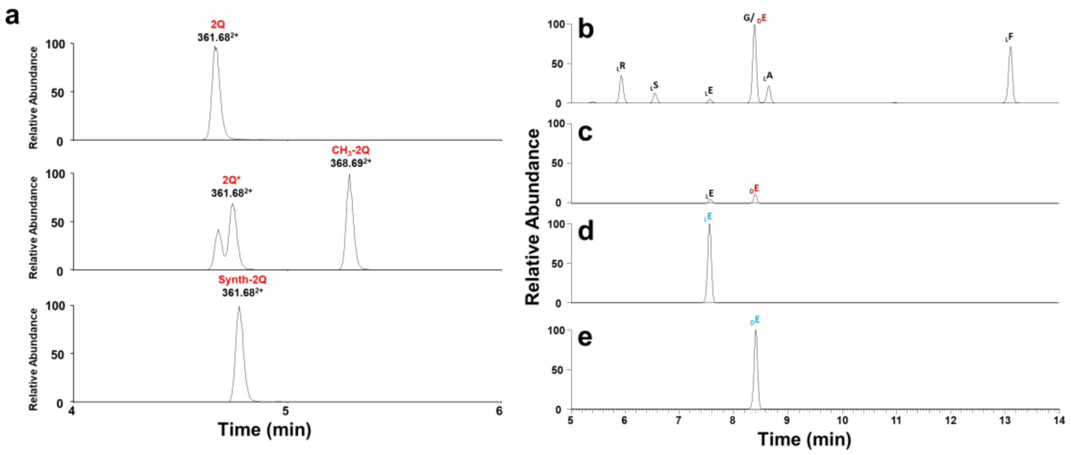

**Figure S17** - LC-MS analysis of peptide **2Q** incubated with **QCMT$_1$**.
(a) Reaction was analyzed at T0 (upper panel) and after 2h (middle panel) and compared with ***synth*-2Q*** (lower panel) containing a D-Glutamine residue in position 5. Incubation was performed under anaerobic conditions in the presence of SAM (0.5 mM), OHCbl (100 µM) and titanium (III) citrate (2 mM). (b) LC-MS analysis of the amino acid content of peptides **2Q** and **2Q*** after acid hydrolysis and L-FDVA derivatization (*see methods*). Of note, Q is converted into E during acidic hydrolysis.[26b] (c) Signal extraction of $_L$E and $_D$E content in the **2Q** and **2Q*** peptides mixture. Reference chromatograms: (d) L-Glu-FDVA and (e) D-Glu-FDVA are indicated by $_L$E and $_D$E, respectively.

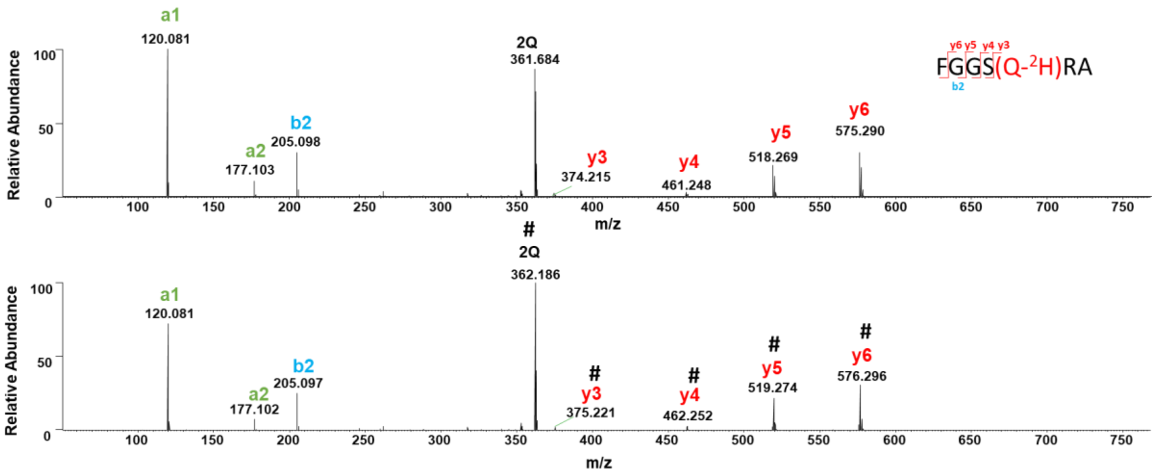

**Figure S18 -** MS-MS fragmentation spectra of **2Q** (upper panel) and **2Q#** (lower panel) in deuterated buffer.

Ions with a mass addition of +1.00627 Da are noted by #.

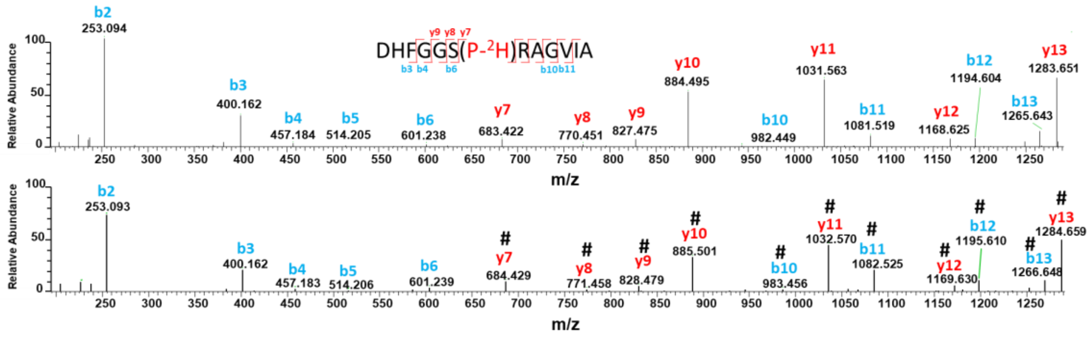

**Figure S19 -** MS-MS fragmentation spectra of **1P** (upper panel) and **1P#** (lower panel) in deuterated buffer. Ions with a mass addition of +1.00627 Da are noted by #.

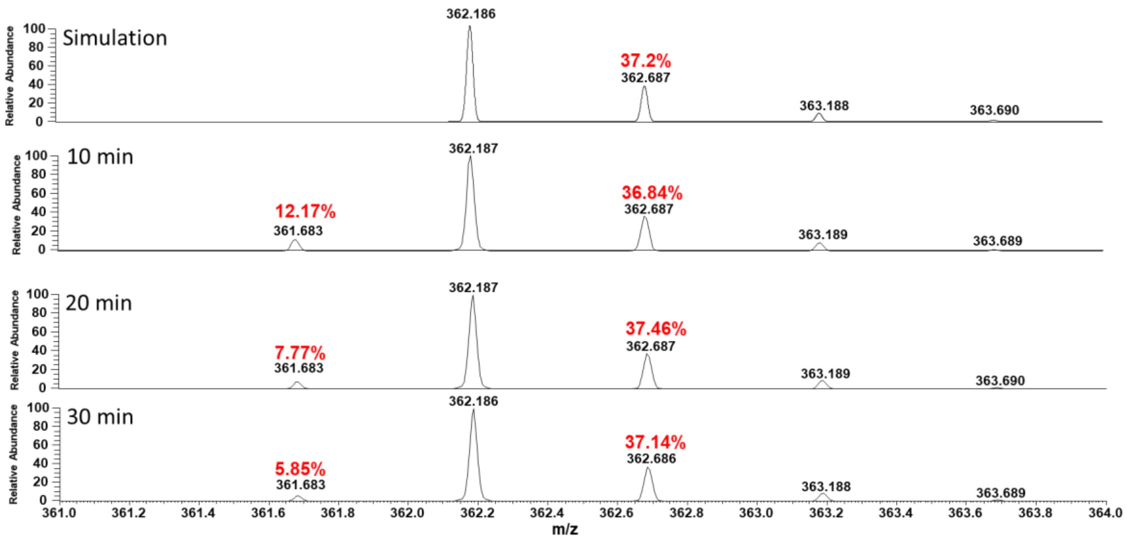

**Figure S20** – Isotopic distribution of **2Q#** over time in deuterated buffer compared to simulated spectrum. The percentages indicated are relative to the first peak of the distribution.

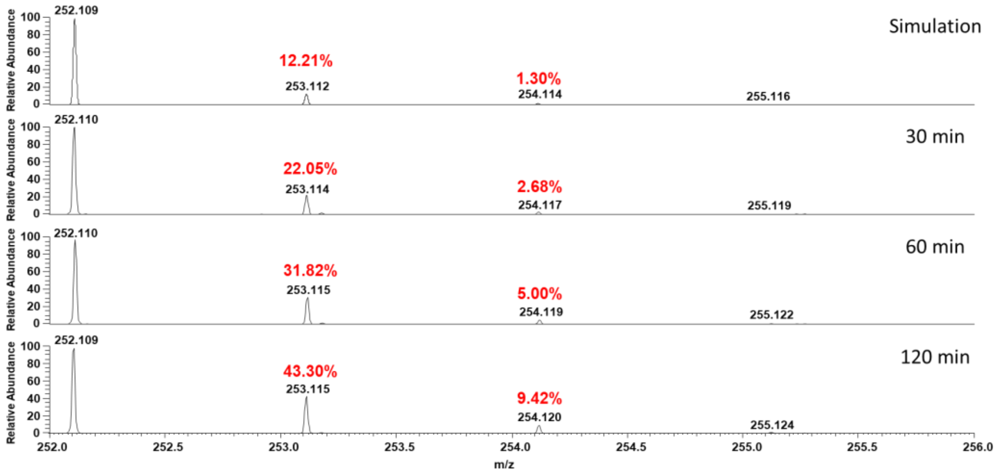

**Figure S21** – Isotopic distribution of **5`dA** over time during reaction with **2Q** in deuterated buffer compared to simulated spectrum.

The percentages indicated are relative to the first peak of the distribution.

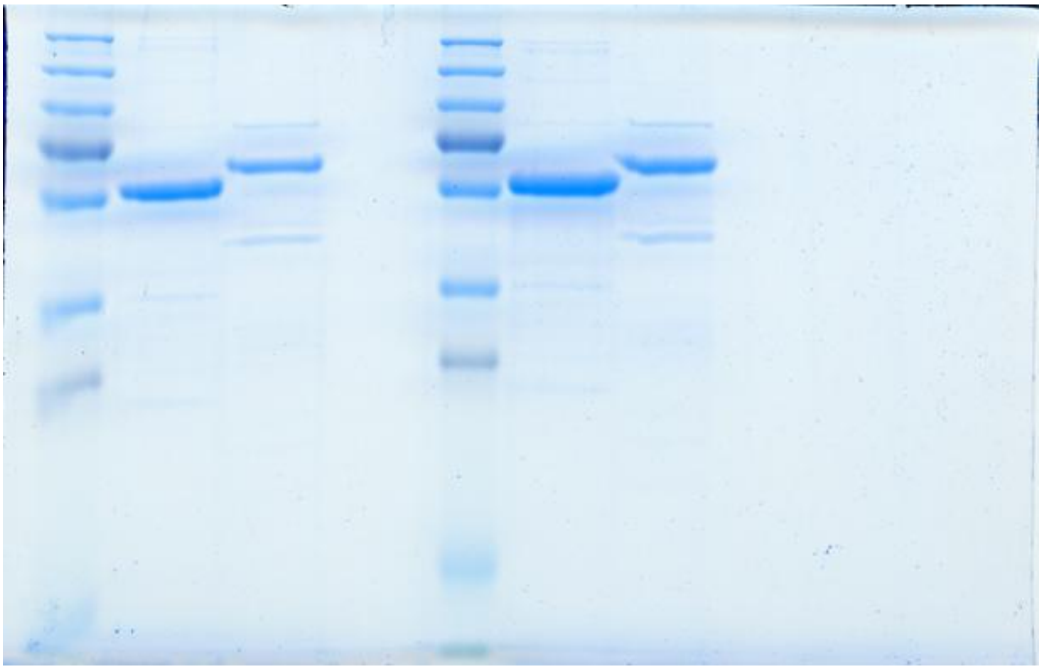

**Figure S22** – SDS-PAGE analysis of **QCMT$_1$** and **QCMT$_2$**.

Uncropped and unedited gel. For annotations, see *Fig. S1*.

| ***a*) Sample details.** | | |
|---|---|---|
| | **$QCMT_1$** | **$QCMT_1$+ OHCob** |
| Organism | *Methanoculleus thermophilus* | |
| UniProt sequence ID (residues in construct) | WP_066957658 | |
| Extinction coefficient [$A_{280}$, 0.1%(*w*/*v*)] | 0.675 | 0.675 |
| Partial specific volume, ν (cm3/g) | 0.732 | 0.732 |
| *M* from chemical composition (kDa) | 50.3 | 51.6 |
| SEC–SAXS column | 5 × 150 mm Superdex increase S200 | |
| Loading concentration (mg $mL^{-1}$) | 5 | |
| Injection volume (µL) | 50 | |
| Flow rate (mL.$min^{-1}$) | 0.3 | |
| Solvent (solvent blanks taken from SEC flowthrough prior to elution of protein) | Tris 50mM, NaCl 300mM, Glycerol 5% DTT 1 mM | |

| ***b*) SAXS data-collection parameters.** | |
|---|---|
| Instrument/data processing | BioSAXS on the SWING beamline at Synchrotron SOLEIL[1] |
| Wavelength (Å) | 1.0332 |
| Beam size (µm) | 500x200 |
| Camera length (m) | 2.00 |
| *q* measurement range ($Å^{-1}$) | 0.0041–0.5516 |
| Absolute scaling method | Comparison with scattering from 1 mm pure $H_2O$ |
| Normalization | To transmitted intensity by beam-stop counter |
| Monitoring for radiation damage | data frame-by-frame comparison |
| Exposure time | Continuous 1 s data-frame measurements of SEC elution |
| Sample configuration | SEC–SAXS with thermalized quartz capillary (ID 1.5mm), Nitrogen gas Buffer and AutoSampler saturated |
| Sample temperature (°C) | 20 |

| **(*c*) Software employed for SAXS data reduction, analysis and interpretation.** | |
|---|---|
| SAXS data reduction | *I*(*q*) *versus q, buffer subtraction & frames selection* using Foxtrot 3.10 (https://www.synchrotron-soleil.fr/en/beamlines/swing#paragraphes_menu_left-block-7) |
| Extinction coefficient estimate | ProtParam (Gasteiger et al., 2005)[2] |
| Basic analysis: Guinier, *P*(*r*), MW | RAW 2.2.1 (Hopkins, 2024, Meisburger et al., 2021, Rambo et al. 2013, Svergun, 1992)[3-6] |
| Three-dimensional graphic model representations | *PyMOL* v.1.70.0.5 Win64 |

| **(*d*) Structural parameters.** | | |
|---|---|---|
| | **$QCMT_1$** | **$QCMT_1$+ OHCob** |
| Guinier analysis | | |
| *I*(0) ($cm^{-1}$) | 0.03 | 0.03 |

| | | |
|---|---|---|
| $R_g$ (Å) | 25.49 | 25.00 |
| $qR_g$ max ($q_{min}$ = 0.0066 $Å^{-1}$) | 1.1 | 1.11 |
| Coefficient of correlation, $R^2$ | 1.0 | 1.0 |
| *MW* from *Vc* (kDa) | 47.2 | 46.1 |
| *P*(*r*) analysis | | |
| *I*(0) ($cm^{-1}$) | 0.03 | 0.03 |
| $R_g$ (Å) | 25.64 | 25.25 |
| $d_{max}$ (Å) | 100 | 100 |
| *q* range ($Å^{-1}$) | 0.01-0.56 | 0.01-0.56 |
| total estimate from *GNOM* | 0.812 | 0.814 |
| **(*f*) Atomistic modelling.** | | |
| Crystal structures | Apo AlphaFold2.0 model aligned with TokK (PDB:7KDX) | |
| *q* range for all modelling | 0.007–0.49 | |
| *PepsiSAXS (*Grudinin et al. 2017)[7] | | |
| $\chi^2$ | 1.00 | 1.10 |
| **(*g*) SASBDB IDs for data and models.** | | |
| | DVY7 | DVZ7 |

**Table S1** - Report of biomolecular structural modelling of small-angle scattering data.

| Peptide | Sequence | Theoretical mass Z = 1 | Theoretical mass Z = 2 | Theoretical mass Z = 3 | Experimental mass | Δppm |
|---|---|---|---|---|---|---|
| **1Q** | DHFGGSQRAGVIA | 1314.655 | 657.831 | 438.890 | 438.891 (Z=3) | 1.8 |
| **1Q*** | DHFGGSQRAGVIA | 1314.655 | 657.831 | 438.890 | 438.891 (Z=3) | 2.6 |
| **1N** | DHFGGSNRAGVIA | 1300.639 | 650.823 | 434.218 | 434.220 (Z=3) | 4.4 |
| **1N*** | DHFGGSNRAGVIA | 1300.639 | 650.823 | 434.218 | 434.219 (Z=3) | 1.4 |
| **1P** | DHFGGSPRAGVIA | 1283.649 | 642.328 | 428.555 | 428.556 (Z=3) | 3.5 |
| **1P*** | DHFGGSPRAGVIA | 1283.649 | 642.328 | 428.555 | 428.556 (Z=3) | 3.6 |
| **1R** | DHFGGSRRAGVIA | 1342.698 | 671.852 | 448.237 | 448.239 (Z=3) | 4.1 |
| **1E** | DHFGGSERAGVIA | 1315.639 | 658.323 | 439.218 | 439.218 (Z=3) | 0.0 |
| **1D** | DHFGGSDRAGVIA | 1301.623 | 651.315 | 434.546 | 434.547 (Z=3) | 2.2 |
| **1Y** | DHFGGSYRAGVIA | 1349.660 | 675.333 | 450.558 | 450.558 (Z=3) | -1.1 |
| **1Y*** | DHFGGSYRAGVIA | 1349.660 | 675.333 | 450.558 | 450.559 (Z=3) | 2.5 |
| **1A** | DHFGGSARAGVIA | 1257.634 | 629.320 | 419.883 | 419.884 (Z=3) | 2.8 |
| **1G** | DHFGGSGRAGVIA | 1243.618 | 622.313 | 415.211 | 415.211 (Z=3) | 1.4 |
| **2**Q | FGGSQRA | 722.358 | 361.683 | 241.458 | 361.683 (Z=2) | 1.9 |
| **2**Q* | FGGSQRA | 722.358 | 361.683 | 241.458 | 361.681 (Z=2) | -4.4 |
| **2**Q# | FGGS(Q-2H)RA | 723.364 | 362.186 | 241.793 | 362.185 (Z=2) | -3.5 |
| **3**Q | FGGSQR | 651.321 | 326.164 | 217.778 | 326.164 (Z=2) | -0.4 |
| **3**Q* | FGGSQR | 651.321 | 326.164 | 217.778 | 326.164 (Z=2) | -1.0 |
| **4**Q | FGGSQ | 495.220 | 248.114 | 165.745 | 495.221 (Z=1) | 3.3 |
| **5**Q | GGSQRA | 575.290 | 288.148 | 192.435 | 288.149 (Z=2) | 1.1 |
| **2A-*d3*** | FGGS(*d*3A)RA | 668.355 | 334.681 | 223.457 | 334.681 (Z=2) | -2.4 |
| **2A-*d4*** | FGGS(*d*4A)RA | 669.362 | 335.184 | 223.792 | 335.184 (Z=2) | -0.1 |

**Table S2 -** Peptide sequences with their theoretical and experimental masses.

*: epimerized peptide. #: deuterated peptide.

| Peptide | Sequence | Theoretical mass Z = 1 | Theoretical mass Z = 2 | Theoretical mass Z = 3 | Experimental mass | Δppm |
|---|---|---|---|---|---|---|
| **$CH_3$-1Q** | DHFGGSQ$^{\Delta}$RAGVIA | 1328.671 | 664.839 | 443.562 | 443.564 (Z=3) | 4.9 |
| **$CH_3$-1N** | DHFGGSN$^{\Delta}$RAGVIA | 1314.655 | 657.831 | 438.890 | 438.891 (Z=3) | 2.8 |
| **$CH_3$-1P** | DHFGGSP$^{\Delta}$RGVIA | 1297.665 | 649.336 | 433.227 | 433.226 (Z=3) | -0.7 |
| **$CH_3$-1R** | DHFGGSR$^{\Delta}$RAGVIA | 1356.713 | 678.860 | 452.909 | 452.911 (Z=3) | 2.8 |
| **$CH_3$-1E** | DHFGGSE$^{\Delta}$RAGVIA | 1329.655 | 665.331 | 443.890 | 443.893 (Z=3) | 6.6 |
| **$CH_3$-1D** | DHFGGSD$^{\Delta}$RAGVIA | 1315.639 | 658.323 | 439.218 | 439.219 (Z=3) | 2.8 |
| **$CH_3$-1Y** | DHFGGSY$^{\Delta}$RAGVIA | 1363.675 | 682.341 | 455.230 | 455.231 (Z=3) | 2.2 |
| **$CH_3$-1A** | DHFGGSA$^{\Delta}$RAAGVIA | 1363.675 | 682.341 | 424.555 | 424.555 (Z=3) | 1.0 |
| **$CH_3$-1G** | DHFGGSG$^{\Delta}$RAAGVIA | 1271.649 | 636.328 | 419.883 | 419.882 (Z=3) | -1.1 |
| **$CH_3$-2Q** | FGGSQ$^{\Delta}$RA | 736.374 | 368.691 | 246.129 | 368.692 (Z=2) | 4.1 |
| **$CH_3$-3Q** | FGGSQ$^{\Delta}$R | 665.337 | 333.172 | 222.450 | 333.172 (Z=2) | 1.2 |
| **$CH_3$-4Q** | FGGSQ$^{\Delta}$ | 509.235 | 255.121 | 170.417 | - | - |
| **$CH_3$-5Q** | GGSQ$^{\Delta}$RA | 589.305 | 295.156 | 197.107 | - | - |
| **$CH_3$-2A-*d3*** | FGGS(d3A)$^{\Delta}$RA | 682.371 | 341.689 | 228.129 | 341.691 (Z=2) | 4.0 |
| **$CH_3$-2A-*d4*** | FGGS(d4A)$^{\Delta}$RA | 682.371 | 341.689 | 228.129 | 341.690 (Z=2) | 2.7 |

**Table S3** - Methylated-peptides with their theoretical and experimental masses.

Positions of the +14.015 Da mass addition, corresponding to an addition of a $CH_2$ are noted by $^{\Delta}$.

| | | 1Q | | $CH_3$-1Q | | |
|---|---|---|---|---|---|---|
| | Sequence | b | y | b | y | |
| 1 | D | 116.034 | 1314.655 | 116.034 | 1328.671 | 13 |
| 2 | H | 253.093 | 1199.628 | 253.093 | 1213.644 | 12 |
| 3 | F | 400.162 | 1062.569 | 400.162 | 1076.585 | 11 |
| 4 | G | 457.183 | 915.501 | 457.183 | 929.516 | 10 |
| 5 | G | 514.205 | 858.479 | 514.205 | 872.495 | 9 |
| 6 | S | 601.237 | 801.458 | 601.237 | 815.473 | 8 |
| 7 | Q | 729.295 | 714.426 | 743.311 | 728.441 | 7 |
| 8 | R | 885.396 | 586.367 | 899.412 | 586.367 | 6 |
| 9 | A | 956.433 | 430.266 | 970.449 | 430.266 | 5 |
| 10 | G | 1013.455 | 359.229 | 1027.470 | 359.229 | 4 |
| 11 | V | 1112.523 | 302.207 | 1126.539 | 302.207 | 3 |
| 12 | I | 1225.607 | 203.139 | 1239.623 | 203.139 | 2 |
| 13 | A | 1296.644 | 90.055 | 1310.660 | 90.055 | 1 |

| | | 1N | | $CH_3$-1N | | |
|---|---|---|---|---|---|---|
| | Sequence | b | y | b | y | |
| 1 | D | 116.034 | 1300.639 | 116.034 | 1314.655 | 13 |
| 2 | H | 253.093 | 1185.612 | 253.093 | 1199.628 | 12 |
| 3 | F | 400.162 | 1048.553 | 400.162 | 1062.569 | 11 |
| 4 | G | 457.183 | 901.485 | 457.183 | 915.501 | 10 |
| 5 | G | 514.205 | 844.464 | 514.205 | 858.479 | 9 |
| 6 | S | 601.237 | 787.442 | 601.237 | 801.458 | 8 |
| 7 | N | 715.279 | 700.410 | 729.295 | 714.426 | 7 |
| 8 | R | 871.381 | 586.367 | 885.396 | 586.367 | 6 |
| 9 | A | 942.418 | 430.266 | 956.433 | 430.266 | 5 |
| 10 | G | 999.439 | 359.229 | 1013.455 | 359.229 | 4 |
| 11 | V | 1098.508 | 302.207 | 1112.523 | 302.207 | 3 |
| 12 | I | 1211.592 | 203.139 | 1225.607 | 203.139 | 2 |
| 13 | A | 1282.629 | 90.055 | 1296.644 | 90.055 | 1 |

| | | 1P | | $CH_3$-1P | | |
|---|---|---|---|---|---|---|
| | Sequence | b | y | b | y | |
| 1 | D | 116.034 | 1283.649 | 116.034 | 1297.665 | 13 |
| 2 | H | 253.093 | 1168.622 | 253.093 | 1182.638 | 12 |
| 3 | F | 400.162 | 1031.563 | 400.162 | 1045.579 | 11 |
| 4 | G | 457.183 | 884.495 | 457.183 | 898.511 | 10 |
| 5 | G | 514.205 | 827.473 | 514.205 | 841.489 | 9 |
| 6 | S | 601.237 | 770.452 | 601.237 | 784.468 | 8 |
| 7 | P | 698.289 | 683.420 | 712.305 | 697.436 | 7 |
| 8 | R | 854.390 | 586.367 | 868.406 | 586.367 | 6 |
| 9 | A | 925.428 | 430.266 | 939.443 | 430.266 | 5 |
| 10 | G | 982.449 | 359.229 | 996.465 | 359.229 | 4 |
| 11 | V | 1081.517 | 302.207 | 1095.533 | 302.207 | 3 |
| 12 | I | 1194.601 | 203.139 | 1208.617 | 203.139 | 2 |
| 13 | A | 1265.639 | 90.055 | 1279.654 | 90.055 | 1 |

| | | 1R | | $CH_3$-1R | | |
|---|---|---|---|---|---|---|
| | Sequence | b | y | b | y | |
| 1 | D | 116.034 | 1342.698 | 116.034 | 1356.713 | 13 |
| 2 | H | 253.093 | 1227.671 | 253.093 | 1241.686 | 12 |
| 3 | F | 400.162 | 1090.612 | 400.162 | 1104.627 | 11 |
| 4 | G | 457.183 | 943.543 | 457.183 | 957.559 | 10 |
| 5 | G | 514.205 | 886.522 | 514.205 | 900.537 | 9 |
| 6 | S | 601.237 | 829.500 | 601.237 | 843.516 | 8 |
| 7 | R | 757.338 | 742.468 | 771.353 | 756.484 | 7 |
| 8 | R | 913.439 | 586.367 | 927.454 | 586.367 | 6 |
| 9 | A | 984.476 | 430.266 | 998.492 | 430.266 | 5 |
| 10 | G | 1041.497 | 359.229 | 1055.513 | 359.229 | 4 |
| 11 | V | 1140.566 | 302.207 | 1154.581 | 302.207 | 3 |
| 12 | I | 1253.650 | 203.139 | 1267.665 | 203.139 | 2 |
| 13 | A | 1324.687 | 90.055 | 1338.703 | 90.055 | 1 |

| | | 1E | | $CH_3$-1E | | |
|---|---|---|---|---|---|---|
| | Sequence | b | y | b | y | |
| 1 | D | 116.034 | 1315.639 | 116.034 | 1329.655 | 13 |
| 2 | H | 253.093 | 1200.612 | 253.093 | 1214.628 | 12 |
| 3 | F | 400.162 | 1063.553 | 400.162 | 1077.569 | 11 |
| 4 | G | 457.183 | 916.485 | 457.183 | 930.500 | 10 |
| 5 | G | 514.205 | 859.463 | 514.205 | 873.479 | 9 |
| 6 | S | 601.237 | 802.442 | 601.237 | 816.457 | 8 |
| 7 | E | 730.279 | 715.410 | 744.295 | 729.425 | 7 |
| 8 | R | 886.380 | 586.367 | 900.396 | 586.367 | 6 |
| 9 | A | 957.417 | 430.266 | 971.433 | 430.266 | 5 |
| 10 | G | 1014.439 | 359.229 | 1028.454 | 359.229 | 4 |
| 11 | V | 1113.507 | 302.207 | 1127.523 | 302.207 | 3 |
| 12 | I | 1226.591 | 203.139 | 1240.607 | 203.139 | 2 |
| 13 | A | 1297.628 | 90.055 | 1311.644 | 90.055 | 1 |

| | | 1D | | $CH_3$-1D | | |
|---|---|---|---|---|---|---|
| | Sequence | b | y | b | y | |
| 1 | D | 116.034 | 1301.623 | 116.034 | 1315.639 | 13 |
| 2 | H | 253.093 | 1186.596 | 253.093 | 1200.612 | 12 |
| 3 | F | 400.162 | 1049.537 | 400.162 | 1063.553 | 11 |
| 4 | G | 457.183 | 902.469 | 457.183 | 916.485 | 10 |
| 5 | G | 514.205 | 845.448 | 514.205 | 859.463 | 9 |
| 6 | S | 601.237 | 788.426 | 601.237 | 802.442 | 8 |
| 7 | D | 716.263 | 701.394 | 730.279 | 715.410 | 7 |
| 8 | R | 872.365 | 586.367 | 886.380 | 586.367 | 6 |
| 9 | A | 943.402 | 430.266 | 957.417 | 430.266 | 5 |
| 10 | G | 1000.423 | 359.229 | 1014.439 | 359.229 | 4 |
| 11 | V | 1099.492 | 302.207 | 1113.507 | 302.207 | 3 |
| 12 | I | 1212.576 | 203.139 | 1226.591 | 203.139 | 2 |
| 13 | A | 1283.613 | 90.055 | 1297.628 | 90.055 | 1 |

| | | 1Y | | $CH_3$-1Y | | |
|---|---|---|---|---|---|---|
| | Sequence | b | y | b | y | |
| 1 | D | 116.034 | 1349.660 | 116.034 | 1363.675 | 13 |
| 2 | H | 253.093 | 1234.633 | 253.093 | 1248.648 | 12 |
| 3 | F | 400.162 | 1097.574 | 400.162 | 1111.590 | 11 |
| 4 | G | 457.183 | 950.505 | 457.183 | 964.521 | 10 |
| 5 | G | 514.205 | 893.484 | 514.205 | 907.500 | 9 |
| 6 | S | 601.237 | 836.463 | 601.237 | 850.478 | 8 |
| 7 | Y | 764.300 | 749.430 | 778.316 | 763.446 | 7 |
| 8 | R | 920.401 | 586.367 | 934.417 | 586.367 | 6 |
| 9 | A | 991.438 | 430.266 | 1005.454 | 430.266 | 5 |
| 10 | G | 1048.460 | 359.229 | 1062.475 | 359.229 | 4 |
| 11 | V | 1147.528 | 302.207 | 1161.544 | 302.207 | 3 |
| 12 | I | 1260.612 | 203.139 | 1274.628 | 203.139 | 2 |
| 13 | A | 1331.649 | 90.055 | 1345.665 | 90.055 | 1 |

| | | 1G | | $CH_3$-1G | | |
|---|---|---|---|---|---|---|
| | Sequence | b | y | b | y | |
| 1 | D | 116.034 | 1243.618 | 116.034 | 1257.634 | 13 |
| 2 | H | 253.093 | 1128.591 | 253.093 | 1142.607 | 12 |
| 3 | F | 400.162 | 991.532 | 400.162 | 1005.548 | 11 |
| 4 | G | 457.183 | 844.464 | 457.183 | 858.479 | 10 |
| 5 | G | 514.205 | 787.442 | 514.205 | 801.458 | 9 |
| 6 | S | 601.237 | 730.421 | 601.237 | 744.436 | 8 |
| 7 | G | 658.258 | 643.389 | 672.274 | 657.404 | 7 |
| 8 | R | 814.359 | 586.367 | 828.375 | 586.367 | 6 |
| 9 | A | 885.396 | 430.266 | 899.412 | 430.266 | 5 |
| 10 | G | 942.418 | 359.229 | 956.433 | 359.229 | 4 |
| 11 | V | 1041.486 | 302.207 | 1055.502 | 302.207 | 3 |
| 12 | I | 1154.570 | 203.139 | 1168.586 | 203.139 | 2 |
| 13 | A | 1225.607 | 90.055 | 1239.623 | 90.055 | 1 |

| | | 1A | | $CH_3$-1A | | |
|---|---|---|---|---|---|---|
| | Sequence | b | y | b | y | |
| 1 | D | 116.034 | 1257.634 | 116.034 | 1271.649 | 13 |
| 2 | H | 253.093 | 1142.607 | 253.093 | 1156.622 | 12 |
| 3 | F | 400.162 | 1005.548 | 400.162 | 1019.563 | 11 |
| 4 | G | 457.183 | 858.479 | 457.183 | 872.495 | 10 |
| 5 | G | 514.205 | 801.458 | 514.205 | 815.473 | 9 |
| 6 | S | 601.237 | 744.436 | 601.237 | 758.452 | 8 |
| 7 | A | 672.274 | 657.404 | 686.289 | 671.420 | 7 |
| 8 | G | 828.375 | 586.367 | 842.390 | 586.367 | 6 |
| 9 | A | 899.412 | 430.266 | 913.428 | 430.266 | 5 |
| 10 | G | 956.433 | 359.229 | 970.449 | 359.229 | 4 |
| 11 | V | 1055.502 | 302.207 | 1069.517 | 302.207 | 3 |
| 12 | I | 1168.586 | 203.139 | 1182.601 | 203.139 | 2 |
| 13 | A | 1239.623 | 90.055 | 1253.639 | 90.055 | 1 |

| | | 2Q | | $CH_3$-2Q | | |
|---|---|---|---|---|---|---|
| | Sequence | b | y | b | y | |
| 1 | F | 148.076 | 722.358 | 148.076 | 736.374 | 7 |
| 2 | G | 205.097 | 575.290 | 205.097 | 589.305 | 6 |
| 3 | G | 262.119 | 518.268 | 262.119 | 532.284 | 5 |
| 4 | S | 349.151 | 461.247 | 349.151 | 475.262 | 4 |
| 5 | Q | 477.209 | 374.215 | 491.225 | 388.230 | 3 |
| 6 | R | 633.310 | 246.156 | 647.326 | 246.156 | 2 |
| 7 | A | 704.347 | 90.055 | 718.363 | 90.055 | 1 |

| | | Synth-2Q | | |
|---|---|---|---|---|
| | Sequence | b | y | |
| 1 | F | 148.076 | 722.358 | 7 |
| 2 | G | 205.097 | 575.290 | 6 |
| 3 | G | 262.119 | 518.268 | 5 |
| 4 | S | 349.151 | 461.247 | 4 |
| 5 | D-Q | 477.209 | 374.215 | 3 |
| 6 | R | 633.310 | 246.156 | 2 |
| 7 | A | 704.347 | 90.055 | 1 |

| | | 3Q | | $CH_3$-3Q | | |
|---|---|---|---|---|---|---|
| | Sequence | b | y | b | y | |
| 1 | F | 148.076 | 651.321 | 148.076 | 665.337 | 6 |
| 2 | G | 205.097 | 504.253 | 205.097 | 518.268 | 5 |
| 3 | G | 262.119 | 447.231 | 262.119 | 461.247 | 4 |
| 4 | S | 349.151 | 390.210 | 349.151 | 404.225 | 3 |
| 5 | Q | 477.209 | 303.178 | 491.225 | 317.193 | 2 |
| 6 | R | 633.310 | 175.119 | 647.326 | 175.119 | 1 |

| | | 4Q | | $CH_3$-4Q | | |
|---|---|---|---|---|---|---|
| | Sequence | b | y | b | y | |
| 1 | F | 148.076 | 495.220 | 148.076 | 509.235 | 5 |
| 2 | G | 205.097 | 348.151 | 205.097 | 362.167 | 4 |
| 3 | G | 262.119 | 291.130 | 262.119 | 305.146 | 3 |
| 4 | S | 349.151 | 234.108 | 349.151 | 248.124 | 2 |
| 5 | Q | 477.209 | 147.076 | 491.225 | 161.092 | 1 |

| | | 5Q | | $CH_3$-5Q | | |
|---|---|---|---|---|---|---|
| | Sequence | b | y | b | y | |
| 1 | G | 58.029 | 575.290 | 58.029 | 589.310 | 6 |
| 2 | G | 115.050 | 518.268 | 115.050 | 532.288 | 5 |
| 3 | S | 202.082 | 461.247 | 202.082 | 475.267 | 4 |
| 4 | Q | 330.141 | 374.215 | 344.161 | 388.235 | 3 |
| 5 | R | 486.242 | 246.156 | 500.262 | 246.156 | 2 |
| 6 | A | 557.279 | 90.055 | 571.299 | 90.055 | 1 |

**Table S4 -** Theoretical ion fragments from peptides used in this study.

Methylated peptides are annotated with “$CH_3$-“.
b and y fragments are numbered respectively in blue (left side) and red (right side).
